# Genome of the Predatory Volute *Melo melo* Provides Insights into Adaptive Gene Family Diversification in a Basal Neogastropod Lineage

**DOI:** 10.1101/2025.10.31.685756

**Authors:** Biswajit Panda, Anik Dey, Rohan Nath, Arunkumar Krishnan

## Abstract

Neogastropods represent one of the most diverse and ecologically specialized lineages of marine mollusks, yet many of their early-diverging families remain genomically underexplored. Here, we present the first high-contiguity genome of a volutid species, the tropical predatory snail Melo melo, assembled using PacBio HiFi long reads. The final assembly spans 2.29 Gb with high contiguity (N50: 18.6 Mb) and completeness (BUSCO: 92.4%). Phylogenomic analyses place M. melo as an early-branching neogastropod lineage, supporting the basal position of Volutidae within the order. Comparative genomic analyses reveal expansions of gene families associated with digestion and nutrient metabolism (e.g., metallopeptidases), neurotoxicity (conotoxin-like genes), neurotransmission (ion channels), detoxification (cytochrome P450s and ABC transporters), and innate immunity (C1q and LRR-containing proteins), among others. These expansions are consistent with M. melo’s predatory behavior and adaptation to sediment-rich benthic habitats. Notably, M. melo lacks nacre-related genes but retains several biomineralization-related gene families previously implicated in shell and pearl formation, offering a genomic basis for investigating its distinctive non-nacreous phenotype. Orthology and CAFE analyses further reveal both conserved and lineage-specific dynamics of gene families across the analyzed gastropods. This study fills a key phylogenetic gap in molluscan genomics and establishes Melo melo as a valuable model for understanding the genomic basis of ecological adaptation and trait evolution in basal neogastropods.

## INTRODUCTION

The Neogastropoda, recognized as one of the most ecologically and morphologically diverse orders of marine gastropods (*1–3*), encompasses over 15,000 extant species distributed globally across a wide range of marine environments (*4*). Noted for their predominantly carnivorous lifestyle, neogastropods exhibit a remarkable array of adaptations, including complex venom systems, specialized feeding structures (e.g., proboscis, radula), and well-developed sensory organs (*5–12*). These traits have facilitated their extensive adaptive radiation (*13*), enabling neogastropods to occupy specialized ecological niches ranging from shallow coastal zones to deep-sea benthic ecosystems (*14–16*).

Although the morphological and ecological diversity of Neogastropoda has been extensively documented (*17–25*), the genomic basis of these adaptations remains relatively underexplored. Despite being a highly speciose clade comprising roughly 60 recognized families (*26*)Whole-genome resources are available for only a few, primarily within the Conidae and Muricidae, leaving significant gaps in our understanding of neogastropod biology. Previous studies on *Conus* and other neogastropod snails have highlighted the role of gene family expansions in driving ecological transitions such as shifts in feeding behavior, venom evolution, and environmental sensing (*7, 27–29*).

Such patterns highlight a broader evolutionary trend in molluscs, especially within gastropods, where the expansion and contraction of gene families play a crucial role in driving the emergence of novel functions and phenotypes (*30*). For example, snails from Panpulmonata and Caenogastropoda lineage show expansion of glycogenin(GYG1), linked to the energy storage needs during aestivation (*31*). The expansion of mucin genes in terrestrial stylommatophorans is thought to underlie their transition from aquatic to terrestrial environments (*32*). Likewise, gastropods inhabiting extreme environments, such as the Neomphalina clade from hydrothermal vents and cold seeps, show extensive expansions in SRCR, HTR4, BTBD6, HSP90, and glutamate regulator gene families, reflecting adaptation to thermal and chemical stress (*33–35*). However, in case of Neogastropods, although earlier studies have mostly focused on venom and toxin genes (e.g., conotoxins and peptidases) (*36–38*), the roles of other functionally important gene families—such as cytochrome P450s, ABC transporters, and leucine-rich repeat (LRR) proteins—remain largely unexplored.

Despite the growing number of genomic studies on Neogastropoda, key questions remain about the origins and evolutionary trajectories of gene family innovations within the clade. Most of the available genomes belong to lineages representing more recently diverged branches of the neogastropod phylogeny (*39–41*), leaving early-diverging families poorly represented. This taxonomic imbalance has constrained broader phylogenomic analyses and limited our understanding of how traits linked to predation and ecological specialization emerged across the group. To bridge these gaps, high-quality genome assemblies are required from basal lineages such as Cancellaridae, Volutidae, and Cystiscidae (*4*), which are essential for reconstructing the tempo and mode of gene family evolution within Neogastropoda.

The family Volutidae, comprising about 83 genera, 12 subfamilies (*26*), is one of the earliest lineages to diverge within Neogastropoda during early Cretaceous (139.8–132.6 mya) (*42*), which displays remarkable ecological and morphological diversity. Members of this family are widely distributed across tropical and temperate oceans, ranging from littoral and shallow sublittoral zones to abyssal depths (*43–45*). Volutids are primarily predatory, inhabiting sandy and muddy substrates where they engulf and narcotize prey using a combination of specialized anatomical adaptations, including an extendable proboscis and biochemically active salivary secretions (*44, 46*). While some species are known to consume bivalves, gastropods, and even echinoderms (*46*), others are specialized predators of predatory gastropods (*47*). Despite their evolutionary significance and ecological breadth, Volutidae remain underrepresented in genomic datasets, with only a single genome—*Fulgoraria chinoi*, generated by short-read sequencing strategies, reported prior to this study.

Here, we present the first high-quality reference genome of *Melo melo* (Lightfoot, 1786), a representative of the family Volutidae, generated using PacBio long-read sequencing. This resource addresses a critical taxonomic and genomic gap in Neogastropoda. Our comparative analyses reveal the expansions of several gene families, including conotoxin-related genes potentially linked to its predatory ecology. Furthermore, phylogenomic analyses using *Melo melo* support the basal placement of Volutidae within Neogastropoda, offering key insights into the early genomic diversification of this ecologically and evolutionarily important molluscan clade.

## MATERIALS AND METHODS

### Sample collection

An adult *Melo melo* individual was collected from the Bay of Bengal (21°26’46.7"N, 87°06’44.3"E) (Chandipur, Odisha, India) **(Figure 1a)**. The snail was transported to the laboratory in seawater. Six distinct tissues (foot, tentacles, siphon, ctenidium, hepatopancreas, and gonads) were dissected out and immediately preserved in RNAlater, then stored at −80 °C until DNA and RNA extraction and subsequent sequencing.

**Figure 1:**
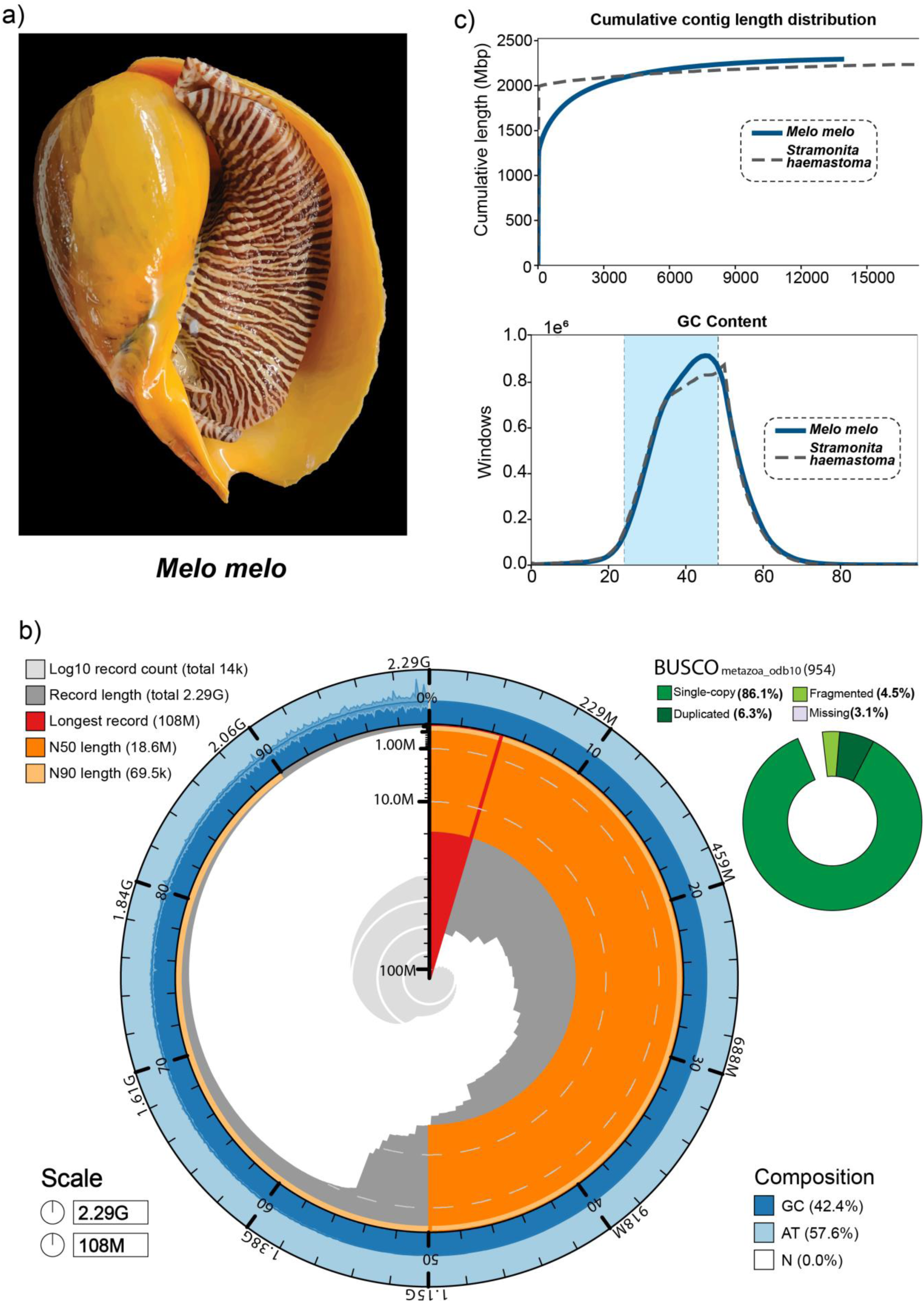
Genome assembly details, completeness and GC content of *Melo melo*. **(a)** Photograph of the sequenced specimen of *Melo melo*. **(b)** BlobToolKit SnailPlot summarizing the final genome assembly of *M. melo*. Circle show total assembly span, longest scaffold, N50, and N90. Inner wedge indicates log10 record count (≈14k scaffolds). Right, BUSCO completeness (metazoa_odb10). **(c)** QUAST summaries. Top: cumulative contig/scaffold length curves showing cumulative length for *M. melo* relative to *S. haemastoma*. Bottom: genome-wide GC-content distribution (42–43%); shaded band denotes the central density for *M. melo*.

### DNA extraction, library preparation and sequencing

High molecular weight genomic DNA was extracted from the foot muscle using a DNA isolation kit (Qiagen) following manufacturer’s instructions. Isolated gDNA quantity was measured using Qubit 4.0 fluorometer (Thermofisher #Q33238, Thermofisher Scientific, Massachusetts, USA) using DNA HS assay kit (Thermofisher #Q32851, Thermofisher Scientific, Massachusetts, USA). DNA purity was checked using NanoDrop 2000 (Thermofisher Scientific, Massachusetts, USA). The integrity of DNA was evaluated on 1% agarose gel and on Agilent FEMTO pulse analyser DNA size was estimated (Agilent Technologies, California, USA). After purification, SMRT library size was prepared, and after size selection was analyzed using Agilent FEMTO pulse analyser (Agilent Technologies, California, USA). The library was constructed using the SMRTbell Express template Preparation Kit 3.0 (Pacific Biosciences, California, USA) as per manufacturers’ protocol. The prepared library was purified using AMPure PB beads (Pacific Biosciences, California, USA). AMPure PB beads were used for size selection to remove fragments shorter than 5 kb. Size. Selected SMRT libraries were purified and then subjected to primer annealing, polymerase binding using Sequel IIe binding kit 3.2 to prepare the bound complex. About 90 pM of the library was loaded on ONE SMRTcell containing 8M ZMW and sequenced in PacBio Sequel IIe system in CCS/HiFi mode.

### *De novo* Genome assembly

The PacBio HiFi reads were *de novo* assembled using Hifiasm (v0.19.9-r616) (*48*), with default parameters. The resulting *de novo* genome assembly was further polished through reference-guided scaffolding, correction, and merging using RagTag (v2.1.0) (*49*). The chromosomal-level genome assembly of *Stramonita haemastoma* (GCA_030674155.1) was considered for this reference-based scaffolding process. Assembly statistics were summarized with BlobToolKit (v4.4.2) (*50*), and a Snail plot was generated to visualize scaffold size distribution, N50/L50, GC content, and BUSCO completeness for the *Melo melo* assembly. In addition to the nuclear genome, we also assembled the complete mitochondrial genome of *Melo melo* using MitoHiFi (*51*) and annotated with MITOS2 (*52*). The circularized genome was visualized through Proksee web server(*53*).

### RNA extraction, library preparation and sequencing

Total RNA was isolated using the TRIzol (Invitrogen) from the above-mentioned six tissues and further purified with the NucleoSpin RNA Cleanup Kit (Macherey-Nagel) following the manufacturer’s protocol. The quantity of isolated RNA was measured using the Qubit 4.0 Fluorometer (Thermo Fisher Scientific) with the Qubit RNA HS Assay Kit (Thermo Fisher Scientific). RNA purity was assessed using a NanoDrop 2000 Spectrophotometer (Thermo Fisher Scientific, Massachusetts, USA). RNA integrity was evaluated via 1% agarose gel electrophoresis, and the RNA Integrity Number (RIN) was determined using the Agilent Bioanalyzer 2100 with the RNA 6000 Nano Kit (Agilent Technologies). For selected high-quality RNA samples, full-length cDNA synthesis was performed using the NEBNext Single Cell/Low Input cDNA Synthesis & Amplification Module (New England Biolabs). After amplification, the full-length cDNA was quantified with the Qubit dsDNA HS Kit and analyzed on a Bioanalyzer 2100 using the High Sensitivity DNA Kit. Library preparation was conducted using the SMRTbell Express Template Preparation Kit 3.0 (PacBio) according to the manufacturer’s instructions. Purified SMRTbell libraries (Pool M2) were evaluated on the FEMTO Pulse System (Agilent Technologies) to determine the final library size distribution. The libraries underwent primer annealing and polymerase binding using the Sequel IIe Binding Kit 3.1 (PacBio) to form the bound complex. Approximately 80 pM of the library was loaded onto a SMRT Cell (8M ZMWs) and sequenced on the PacBio Sequel IIe system in CCS/HiFi mode. The raw Iso-Seq reads were processed using the PacBio Iso-Seq v4.0.0 pipeline with default parameters to generate high-quality isoforms, which were subsequently used for genome annotation.

### Assembly Quality Assessment

To evaluate the completeness and overall quality of the *Melo melo* genome assembly, we employed multiple complementary tools. BUSCO (Benchmarking Universal Single-Copy Orthologs, v5.7.1) (*54*) was used to assess genome completeness based on the presence of conserved orthologous genes from the metazoa_odb10 lineage dataset. QUAST (Quality Assessment Tool for Genome Assemblies, v5.3.0) (*55*) was utilized to calculate key assembly statistics including N50, L50, total length, and GC content. To examine read quality and assess for possible contaminants or sequencing artifacts, FastQC (v0.11.9) (*56*) was run on the raw PacBio HiFi reads.

### Repeat Identification, Gene Prediction, and Functional Annotation

Repeat identification in the *Melo melo* genome was performed using both homology-based and *de-novo* approaches. Homology-based repeat masking was carried out using RepeatMasker v4.1.7-p1 with Repbase libraries (*57*). *De-novo* repeat discovery and modeling were conducted using RepeatModeler v2.0.6 (*58*). Gene prediction was performed using BRAKER2 v2.1.6 (*59*), which combines *ab-initio* and transcriptome-guided approaches. Full-length Iso-Seq transcripts from *M. melo* were aligned to repeat-masked genome assemblies using Minimap2 v2.17 (*60*) with splice-aware settings (-ax splice). The resulting alignments were provided to BRAKER2 as extrinsic evidence. GeneMark-ETP (*61*) and AUGUSTUS were trained on these alignments to produce high-confidence gene models. Functional annotation of the predicted protein-coding genes was performed using a multi-database strategy to assign putative functions, domains, and pathway associations. Redundant protein sequences were clustered using CD-HIT v4.8.1 (*62*) with a 95% identity threshold, and sequences shorter than 150 amino acids were excluded to remove likely truncated predictions. The filtered proteins were then annotated using InterProScan v5.59-91.0 (integrating databases such as Pfam, SMART, and PANTHER), Swiss-Prot (UniProtKB 2023_03 release) via BLASTp (BLAST+ v2.13.0), and KEGG pathways using KAAS. An e-value cutoff of 1e−5 was applied in all similarity searches to ensure high-confidence functional assignments.

### Conotoxin gene prediction

Reference conotoxin sequences were retrieved from prior studies on neogastropod venom repertoires (*41, 63, 64*) and from Pfam domain annotations (e.g., PF02950). These were used as queries in a *tblastn* search against the *M. melo* genome. Putative hits were further analyzed using HMMER v3.3 (*65*) with an e-value threshold of 1e−5. Candidate sequences were then validated using HHpred and cross-checked against the ConoServer database (*66*). High-confidence matches were classified as putative conotoxin genes.

### Orthology and Phylogenomic Analysis

Protein-coding genes of *Melo melo* and publicly available transcriptomic and genomic datasets from 21 other molluscan species were retrieved from NCBI and other public repositories **(Supplementary Table 3)**. The dataset included 17 neogastropods spanning four major superfamilies (Conoidea, Buccinoidea, Muricoidea, and Volutoidea), and five outgroup species representing Architaenioglossa, Littorinimorpha, and Heterobranchia. For transcriptome-based taxa, raw reads were quality filtered and adapter-trimmed using standard protocols (e.g., Trimmomatic or fastp), and assembled *de novo* using Trinity (*67*) with default parameters. Gene models were predicted as required, and all sequence headers were standardized prior to phylogenomic analysis. Orthogroup inference followed previously described pipelines (*68, 69*), using OrthoFinder v2.5.4 (*70*) with MCL inflation set to 2.1 which yields homologous sequences in the Orthogroup Sequences folder. These Orthogroups were filtered by removing the redundant sequences shorter than 100 amino acids and using <u>uniqHaplo.pl</u> and were aligned using MAFFT v7.520 (*71*) with the flags –auto and –maxiterate of 1000. Following this low quality and misaligned sequences were filtered out using HmmCleaner (*72*) and BMGE v.1.12 (*73*) was employed to remove regions which were ambiguously aligned. Only those Orthogroups which represented over 75% of the taxa were considered for further downstream analysis. Individual gene trees for each orthogroup were constructed for each using FastTreeMP with gamma rate variation and - slow parameters. Paralog Pruning and exogenous contamination exclusion was done using PhyloPyPruner (*74*). Parameters like: --min-support 0.9, --mask pdist, --trim-lb 3, --trim-divergent 0.75, --min-pdist 0.01 and--prune LS were used. PhyloPyPruner also provided a supermatrix alignment, which was used for phylogenetic analysis. For the phylogenetic tree estimation using Maximum Likelihood (ML), IQ-TREE2 (*75*) was implemented with -T AUTO, -B 1000, --wbtl, model selection set as MPF and 1000 ultrafast bootstrap replicates as the parameters ensuring robust resolution of *Melo melo*’s phylogenetic position within Neogastropoda. Orthology analyses were inferred with OrthoFinder (v2.5.4) across a ten-species panel—five neogastropods, one nudibranch, one aplysiid, one littorinimorph, and two architaenioglossans. Orthogroup intersections are summarized with an UpSet plot, and a focused comparison among the five neogastropods is visualized as a Venn diagram.

### Gene Family Expansion and Contraction

To investigate lineage-specific expansions of gene families in *Melo melo*, we performed a comparative analysis of gene family repertoires across other gastropod genomes in the study. This approach enabled the identification of expansions in gene families associated with key biological processes such as detoxification, immune response, metabolism, and cellular functions—traits often linked to ecological adaptation. Based on functional annotations, gene families were grouped by biological roles, including metabolism (e.g., peptidases), neuronal channels, detoxification (e.g., cytochrome P450s, ABC transporters), and immune response (e.g., C1q and LRRs). To validate these expansions, multiple sequence alignments were generated using MAFFT, and conserved domains were confirmed by comparing with reference sequences from the Pfam database. To strengthen orthology assignments and support observed expansion patterns, gene families were further evaluated through clustering using OrthoFinder.

## RESULTS AND DISCUSSION

### *De Novo* Genome Assembly with Reference-Guided Scaffolding

The PacBio whole genome sequencing resulted in a total 25,103,218,299 bases representing 2852782 raw reads with average read length of 8799 bp **(Supplementary Table1a)**. The initial *de-novo* assembly of *Melo melo* genome using Hifiasm assembler yielded 18,019 contigs, with an N50 of 427.36 Kb and an L50 of 1,441. To further improve the contiguity of the genome, we used a reference-guided scaffolding approach by considering the *Stramonita haemastoma* genome (GCA_030674155.1). This resulted in a more contiguous assembly of 13,961 scaffolds, significantly improving the N50 to 18.6 Mb and reducing the L50 to 27 **(Figure 1b**, **Table 1a, Supplementary Table 1c)**. The final genome assembly size was estimated to be 2.29 Gb for *M. melo*, consistent with the relatively large genome sizes reported in other neogastropods **(Figure 1b)**. Cumulative contig length analysis via QUAST (*55*) showed that *M. melo* has longer contigs early in the assembly index, with cumulative lengths surpassing those of *S. haemastoma*, indicating effective representation of the genomic content **(Figure 1c)**. The assembly exhibits a GC content of 42.45%, and only 0.02% of the genome comprises unresolved gap sequences (“N”), indicating high contiguity and minimal fragmentation **(Figure 1c)**. Genome completeness assessed using BUSCO (metazoa_odb10 dataset) showed 92.4% complete orthologs, including 86.1% single-copy and 6.3% duplicated genes (**Figure 1b**, **Table 1b**). The remaining fragmented and missing orthologs are likely due to the inherent complexity and repeat-rich nature of molluscan genomes. Together, these metrics suggest the high quality and completeness of *M. melo* genome assembly. The complete circular mitochondrial genome of *Melo melo* was recovered, with a total length of 15,827 bp. It contains the expected set of 37 genes, including 13 protein-coding genes (PCGs), 22 transfer RNA (tRNA) genes, and 2 ribosomal RNA (rRNA) genes. The gene order begins with cox3 and ends with trnF. A circular map of the mitogenome is provided in **Supplementary Figure 2**.

**Table 1:** The genome assembly and BUSCO statistics of *Melo melo.* a) Highlights the genome size, number of scaffolds, N50, L50 and GC percentage. b) BUSCO statistics representing the percentage of genome completeness.

| Genome assembly Statistics |  |
| --- | --- |
| Total size of scaffolds | 2294228763 bp |
| Longest scaffold | 107503566 bp |
| Shortest scaffold | 2089 bp |
| Number of scaffolds | 13961 |
| Mean scaffold size | 164331 bp |
| N50 scaffold length | 18608260 bp |
| L50 scaffold count | 27 |
| GC (%) | 42.45 |

| BUSCO Statistics |  |
| --- | --- |
| **C:92.4%[S:86.1%,D:6.3%],F:3.1%,M:4.5%,n:954,E:5.7%** |  |
| Complete BUSCOs (C) | 881 |
| Complete and single-copy BUSCOs (S) | 821 |
| Complete and duplicated BUSCOs (D) | 60 |
| Fragmented BUSCOs (F) | 30 |
| Missing BUSCOs (M) | 43 |
| Total BUSCO groups searched | 954 |

### Repeat Content Analysis

Using RepeatModeler and RepeatMasker, a total of 1,394,082,595 bp of repetitive elements were identified from the genome assembly of *M. melo,* constituting more than half (60.76%) of the assembled genome **(Supplementary Figure 1, Supplementary Table 1d),** aligning with the trend seen among neogastropods (*39, 40, 76*). Of these repetitive elements, transposable elements (TE) were most notable, accounting for 43.05% of them, including long interspersed nuclear elements (LINEs), DNA transposons, long-terminal repeat elements, and some unknown TEs. Furthermore, the Unclassified TEs (which represent 27.50% of the assembled genome) are likely the lineage-specific variants of known repetitive elements. Additionally, LINEs were found to be most prevalent, making up 14.81% of the genome, followed by DNA transposons (0.57%) and LTRs (0.18%) **(Supplementary Table 1d)**. The proliferation of repetitive elements observed in the *M. melo* genome covers a larger proportion of its genome.

Repetitive elements are increasingly recognized as dynamic components of the genome, influencing gene expression, chromatin organization, and overall genomic architecture, while also shaping evolutionary trajectories by promoting genetic variation, structural rearrangements, and regulatory innovation (*77*). In *M. melo*, the substantial proportion of repeats may also partially explain its relatively large genome size, potentially facilitating lineage-specific innovations and ecological specialization. Such TE expansions, frequently observed across gastropod genomes (*78, 79*), have been proposed as adaptive responses to environmental pressures—an interpretation that may also apply to the repeat landscape observed in *M. melo*. In addition, the Kimura divergence profile of *M. melo* reveals a genome dominated by ancient transposable elements **(Figure 2)**. Higher K values indicated a more ancient transposition and vice versa. The TE landscape of *M. melo* indicates a major TE expansion event in the distant past, with little evidence of recent TE activity.

**Figure 2.**
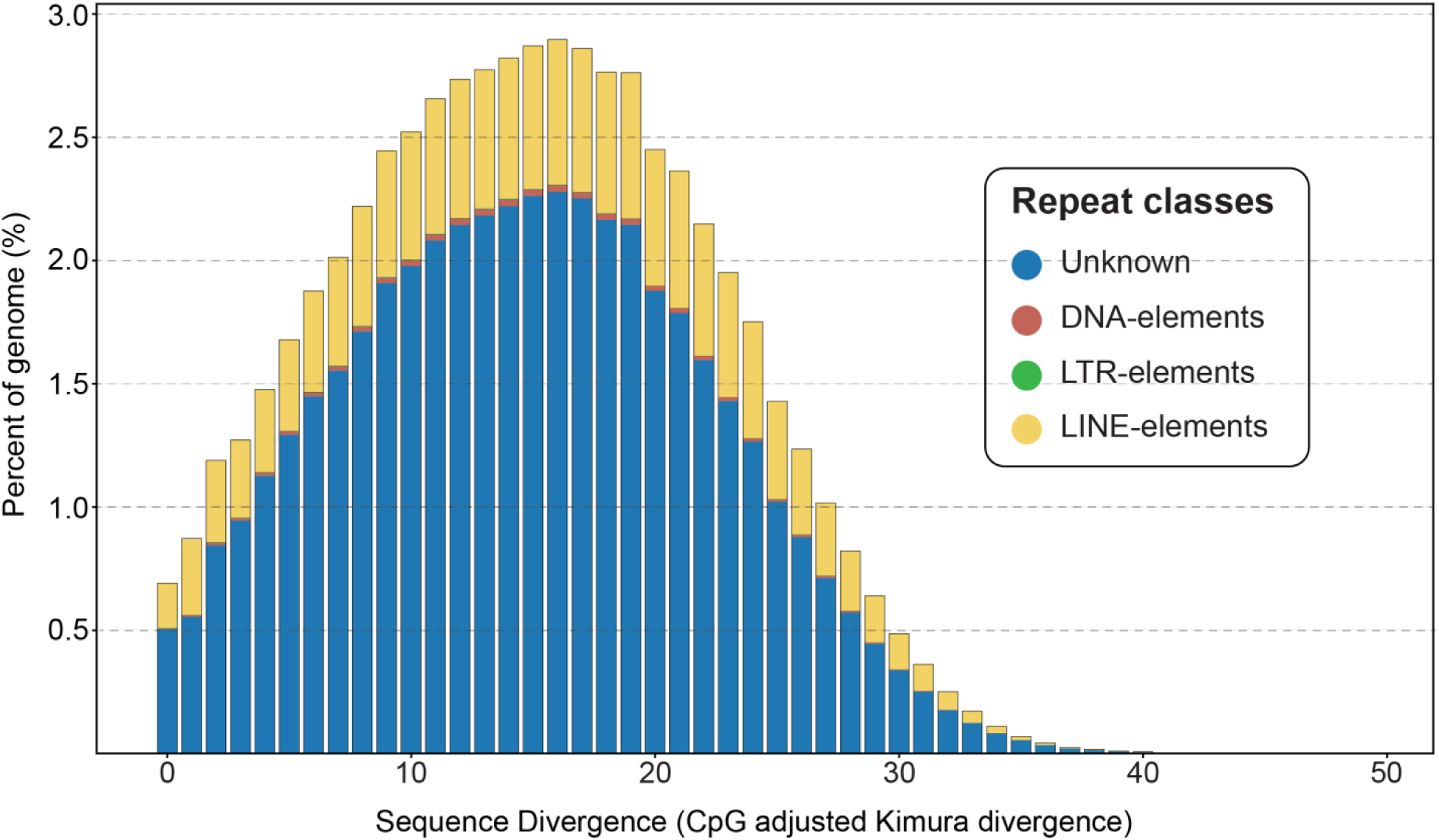
Repeat landscape of *Melo melo* based on CpG-adjusted Kimura divergence. Stacked bars show the proportion of the genome contributed by annotated repeats across binned sequence divergence (x-axis; lower K = more recent, higher K = older). Colors denote repeat classes: blue = Unknown, brown = DNA transposons (DNA-elements), green = LTR retrotransposons (LTR-elements), yellow = LINE retrotransposons (LINE-elements). Y-axis represents the percent of the genome per 1% divergence bin. Repeat divergence was estimated using CpG-adjusted Kimura distances computed by RepeatMasker.

### Gene Prediction and Functional Annotation

A total of 29,364 protein-coding genes were predicted in the *M. melo* genome using BRAKER2, guided by full-length Iso-Seq alignments **(Supplementary Table 2)**. Of these, 26,502 genes were annotated with Pfam domains, reflecting widespread conservation of protein families. SwissProt annotated 24,949 genes, while InterProScan provided annotations for 21,055 genes. KEGG pathway mapping assigned functions to 10,804 genes, indicating substantial representation of core metabolic and signaling pathways **(Supplementary Table 2)**. Both the number of gene predictions provided by BRAKER2 and the number of functionally annotated genes are in agreement with the values observed in Mollusca (*80*).

### Phylogenomics and Early Divergence of *Melo melo* within Neogastropoda

To resolve the evolutionary position of *Melo melo*, we conducted a phylogenomic analysis using nuclear gene alignments from 22 molluscan species spanning 13 gastropod families **(Supplementary Table 3)**. The dataset included key representatives of Neogastropoda, alongside outgroups from Littorinidae, Ampullariidae, Aplysiidae, and Aeolidiidae. A total of 4,718 orthogroups, identified using OrthoFinder, were used to construct the phylogenetic tree. The resulting maximum likelihood tree placed *Melo melo* as the earliest-diverging lineage within Neogastropoda, with strong bootstrap support (>90%), positioning it as sister to all remaining neogastropod taxa **(Figure 3)**. This supports a deep-branching origin for Volutidae and reinforces prior evidence that this family occupies a basal position within neogastropod radiation.

**Figure 3.**
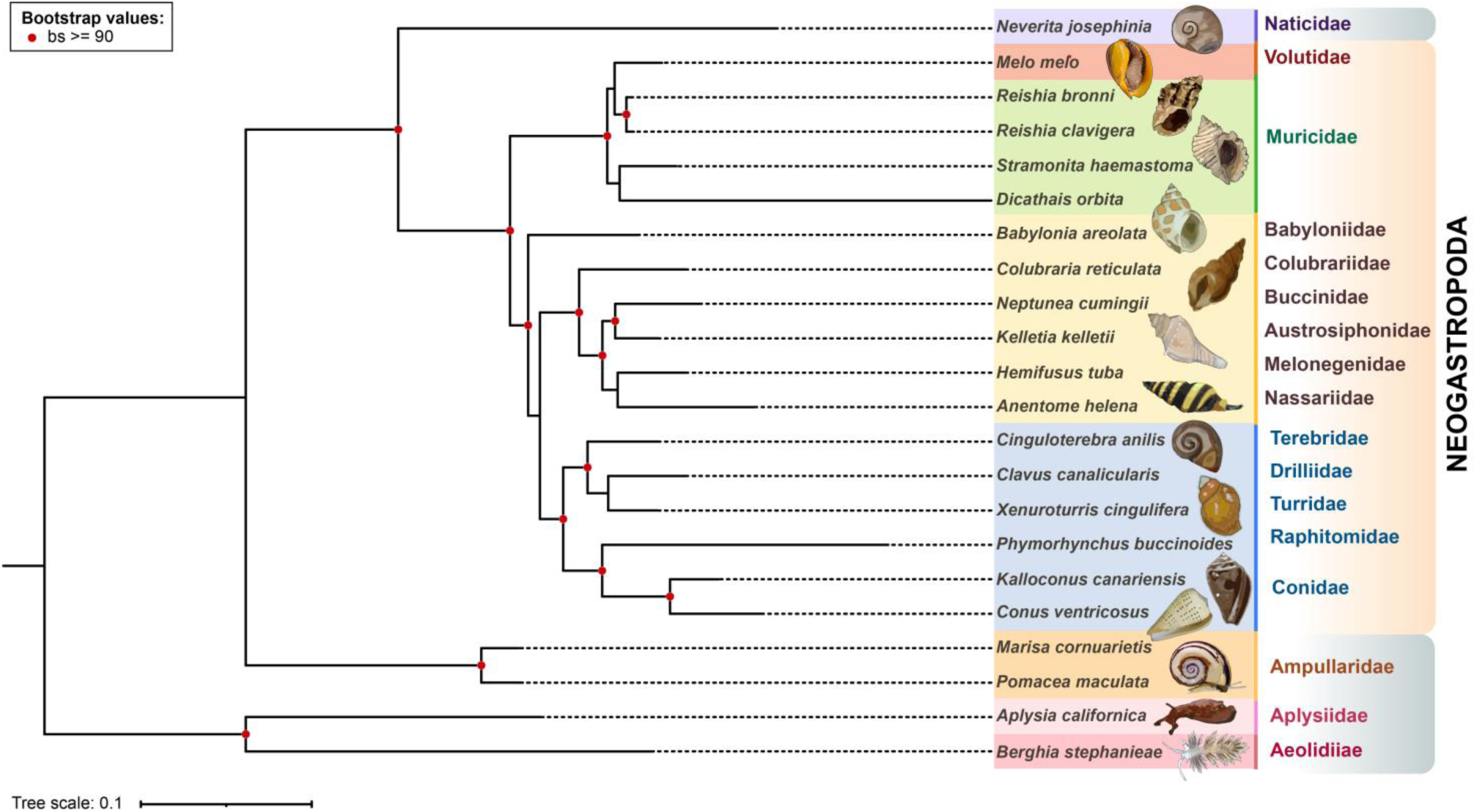
Maximum likelihood phylogeny of *Melo melo* and related neogastropods. Phylogenomic reconstruction based on 4,718 nuclear orthologs across 22 gastropod species, representing 13 families. *Melo melo* (Volutidae) recovered as the earliest-diverging lineage within Neogastropoda, forming a strongly supported clade (bootstrap ≥ 90%) sister to all other sampled neogastropods. Conoidean families (Conidae, Turridae, Drilliidae, Raphitomidae, Terebridae) formed a monophyletic group, while Muricidae, Buccinoidea, and other families showed stable internal relationships. Outgroup taxa from Littorinimorpha, Heterobranchia, and Architaenioglossa were used to root the tree. Colored clades indicate family-level classification. Red circles mark nodes with ≥90% bootstrap support. Scale bar denotes substitutions per site.

Although *Fulgoraria chinoi*—a volutid species previously reported as basal within the family(*81*) —is the only other sequenced member of Volutidae to date, we excluded it from our analysis due to its low genome completeness (21.2% BUSCO) and lack of gene coverage suitable for phylogenomic inference. Nonetheless, mitochondrial studies also support the early divergence of *Melo melo* and other Amoriinae representatives within Volutidae (*43*), suggesting consistent signals across both nuclear and mitochondrial markers. Furthermore, recent molecular phylogenies (*82, 83*) place Volutidae as a distinct lineage separate from Cancellariidae, challenging earlier notions of a shared Volutoidea grouping and underscoring the need to reevaluate these deeper relationships.

Within our tree, all sampled families of Conoidea—Conidae, Turridae, Terebridae, Drilliidae, and Raphitomidae—formed a strongly supported monophyletic group (>90% bootstrap). *Conus ventricosus* and *Kalloconus canariensis* clustered together, while *Xenuroturris cingulifera* (Turridae), *Cinguloterebra anilis* (Terebridae), and *Clavus canalicularis* (Drilliidae) branched successively within the clade **(Figure 3)**. *Phymorhynchus buccinoides* (Raphitomidae) emerged as a terminal lineage within this assemblage, consistent with prior molecular studies of deep-sea conoideans (*4, 84*).

In the buccinoid assemblage, *Babylonia areolata* (Buccinidae) grouped with *Colubraria reticulata* (Colubrariidae), while two additional well-supported clades were recovered: one uniting *Neptunea cumingii* (Buccinidae) and *Kelletia kelletii* (Austrosiphonidae), and another joining *Hemifusus tuba* (Melongenidae) with *Anentome helena* (Nassariidae). Muricidae formed a distinct cluster comprising *Stramonita haemastoma*, *Dicathais orbita*, and *Reishia spp*., reflecting stable family-level relationships **(Figure 3)** previously reported in morphological and molecular studies (*85*). The outgroup taxa—*Neverita josephinia* (Littorinidae), *Pomacea maculata* and *Marisa cornuarietis* (Ampullariidae), *Aplysia californica* (Aplysiidae), and *Berghia stephanieae* (Aeolidiidae)—were placed outside the neogastropod clade, effectively rooting the tree **(Figure 3)** and contextualizing the early diversification of Caenogastropoda (*86*).

While our analysis resolves many family-level relationships with high confidence, it reflects the current lack of genome-scale data for lineages such as Cancellariidae, Olividae, and Mitridae, which could not be included in this reconstruction. Historical classifications grouped Volutidae with several of these families under Volutoidea, but recent phylogenies have shown limited support for this assemblage, often placing Volutidae and Cancellariidae on separate early branches (*43, 82*). The basal position of *Melo melo*—supported by nuclear gene alignments and a high-quality genome—aligns with divergence time estimates for early neogastropod radiation (*87*) (∼132 Ma). This time frame overlaps with the Cenomanian–Turonian anoxic event (89–94 Ma), a period associated with extensive marine ecological turnover. The strong clade-level resolution for Conoidea, Buccinoidea, and Muricidae further reflects the early diversification of carnivorous neogastropods, underscoring the importance of Volutidae in understanding the origins of this major lineage.

### Comparative Genomics and Gene Family Evolution

#### Orthogroup Sharing and Evolutionary Conservation of Gene Families

Orthology inference across 10 representative gastropod species revealed that *Melo melo* retains a large number of conserved orthogroups, while also harboring a distinct set of gene families absent in all other taxa analyzed. A total of 436 orthogroups were unique to *Melo melo*, with no detectable homologs in the remaining species **(Figure 4b)**, suggesting the presence of uncharacterized expansions or lineage-specific innovations in Volutidae. Functional domain analysis of these *Melo*-specific orthogroups revealed prominent enrichment of 7tm_1 (n = 60) and 7TM_GPCR_Srw (n = 19) domains, both associated with G protein-coupled receptors. Other notable domain families included Ldl_recept_a (n = 22), Ank_2 (n = 28), C1q (n = 8), Trypsin (n = 8), LRR_8 (n = 8), and lectin-related modules (e.g., Lectin_C, PAN_1), many of which are involved in immunity, proteolysis, or signal transduction—key features of predatory and immune- adapted mollusks

**Figure 4.**
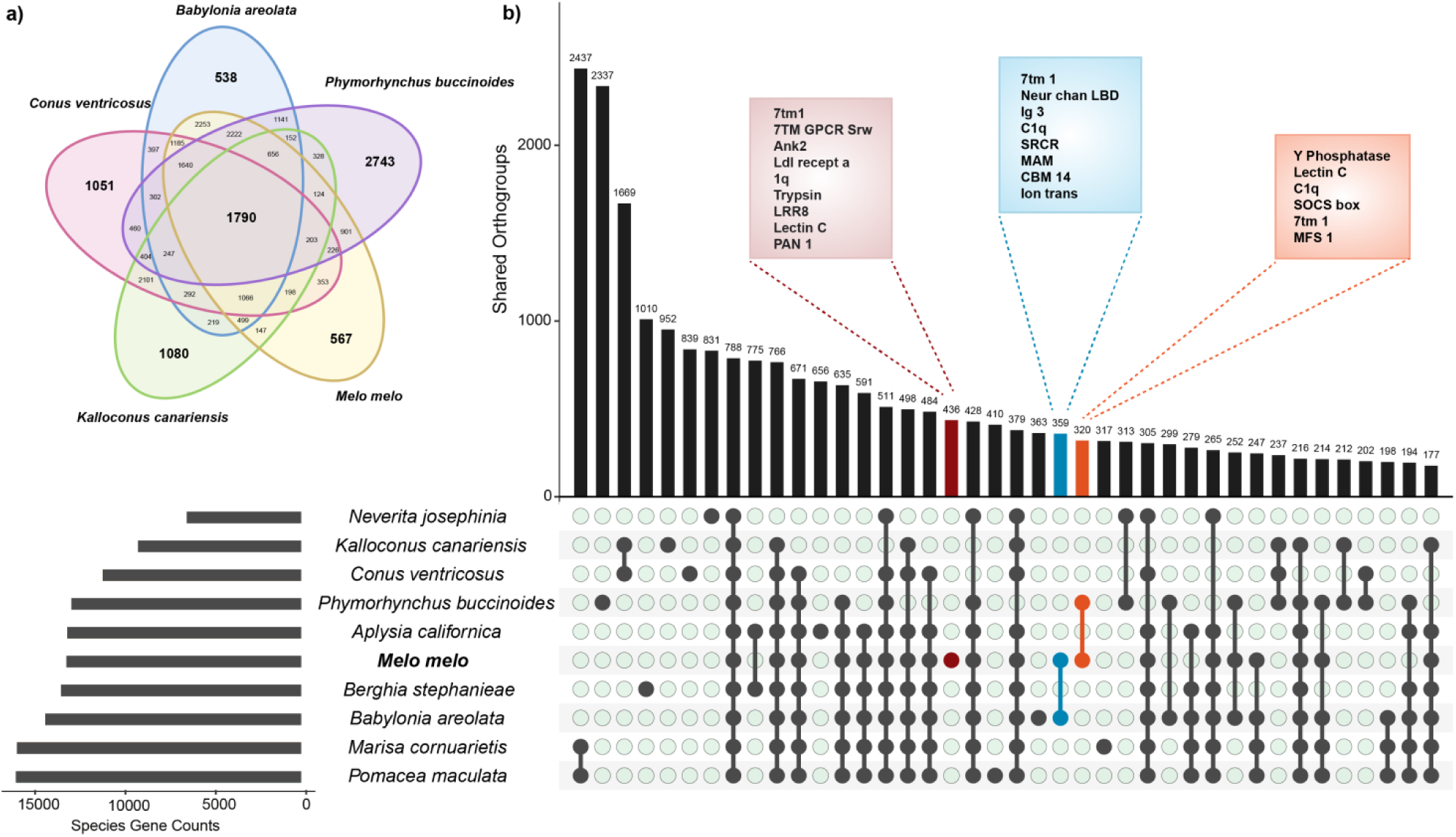
Orthogroup overlap and *Melo melo*-specific gene family expansions across gastropods. **(a)** Venn diagram showing orthogroup sharing among five neogastropod species. **(b)** UpSet plot summarizing orthogroup intersections across ten gastropod species. The orthogroups unique to *Melo* have been highlighted as brick red, in addition the shared orthogroups with *Babylonia areolata* and *Phymorhynchus buccinoides* are also highted as light blue and light orange boxes respectively.

When the analysis was restricted to five neogastropods (*Melo melo*, *Babylonia areolata*, *Phymorhynchus buccinoides*, *Conus ventricosus*, and *Kalloconus canariensis*), *Melo melo* still retained 567 orthogroups with no detectable orthologs in the remaining four taxa **(Figure 4a)**. Several of the same domain families reappeared in this neogastropod-specific comparison, at even higher counts—for instance, 7tm_1 (n = 75), 7TM_GPCR_Srw (n = 30), Ank_2 (n = 26), C1q (n = 20), and Trypsin (n = 15). Additionally, the presence of Integrase_H2C2 and DDE_Tnp_1_7 domains among *Melo*-specific orthogroups hints at possible contributions from transposon-derived expansions, consistent with the high repetitive content observed in the *Melo melo* genome.

Despite this distinctiveness, *Melo melo* also showed considerable overlap with other neogastropods. It shared 359 orthogroups with *Babylonia areolata* and 320 with the deep-sea taxon *Phymorhynchus buccinoides*, with 252 orthogroups commonly retained across all three species **(Figure 4b)**. The shared domains in these overlapping orthogroups included additional sensory and immune-related families such as 7tm_1, Neur_chan_memb, Neur_chan_LBD, Ig_3, C1q, Lectin_C. The strong overlap between *Melo*, *Babylonia*, and *Phymorhynchus*—despite their ecological divergence (coastal vs deep-sea habitats) (*35, 39, 84*) suggests a deeply conserved neogastropod core repertoire, alongside lineage-specific elaborations.

At broader phylogenetic scales, the UpSet analysis suggested that *Melo melo* shares 788 orthogroups with all other gastropods analyzed in the dataset, including *Neverita josephinia*, *Aplysia californica*, and *Berghia stephanieae*. Excluding *Neverita* slightly reduced this set to 766, indicating that the majority of conserved orthogroups in *Melo melo* are broadly retained across caenogastropod and heterobranch lineages. A nested subset of seven taxa, spanning both marine and freshwater gastropods, shared 635 orthogroups, suggesting deep conservation of gene families likely involved in fundamental cellular or physiological processes **(Figure 4b)**. The largest number of shared orthogroups (2437) was observed between the two ampullariid freshwater snails *Pomacea maculata* and *Marisa cornuarietis* **(Figure 4b)**, consistent with their close evolutionary relationship and ecological similarity. Taken together, these orthology patterns underscore the duality of *Melo melo*’s genome: it retains a wide array of evolutionarily conserved gene families while also evolving a unique complement of expansions—particularly in chemosensory, immune, and transposon-related domains—that may underlie the distinct biology of Volutidae.

### Gene Family Evolution in *Melo melo*

Gene family evolution in *Melo melo* was assessed using CAFE5 (*88*), based on orthogroup counts across ten gastropod genomes. Along the lineage leading to *M. melo* (Volutidae), 2443 gene families were inferred to have expanded, while 219 showed contractions **(Figure 5)**. This represents one of the most substantial signatures of gene expansion across the analysed taxa. These expansions may correspond to lineage-specific innovations within Volutidae.

**Figure 5.**
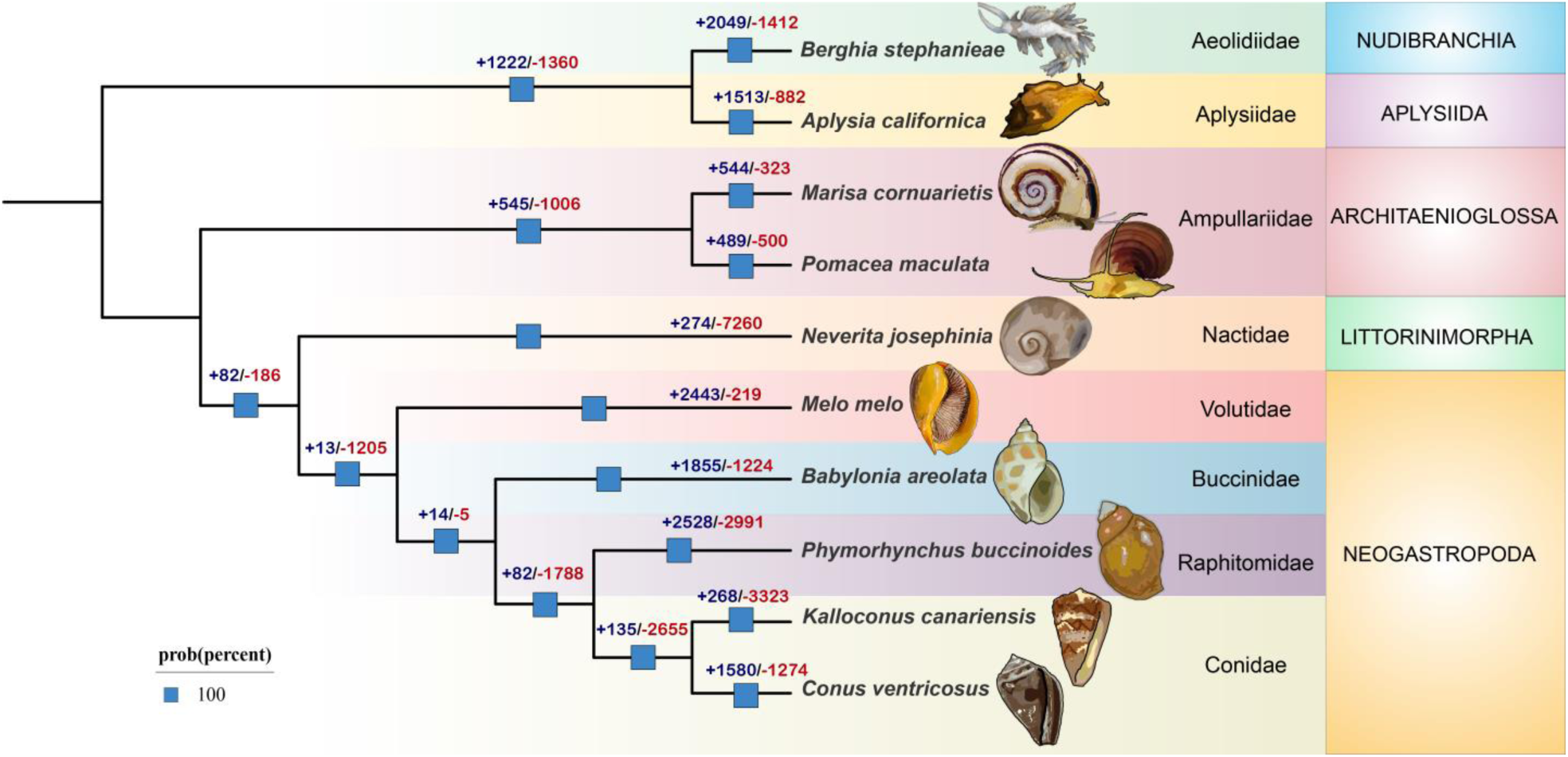
Gene family expansions and contractions across gastropod lineages. CAFE-based analysis of 10 gastropod species reveals extensive gene family turnover across major lineages. Values at internal nodes indicate total gene gains (+, blue) and losses (–, red). All nodes shown had 100% posterior probability.

Other neogastropod taxa also showed notable shifts in gene family size. *Babylonia areolata* (Buccinidae) exhibited 1,855 expansions and 1224 contractions, while *Phymorhynchus buccinoides* (Raphitomidae) displayed 2528 expansions alongside 2991 contractions. The two conid species, *Conus ventricosus* and *Kalloconus canariensis*, showed more moderate expansion counts (1580 and 268, respectively), though the latter experienced substantial gene loss (3323) **(Figure 5)**. Among non-neogastropods, the gene gain and loss patterns varied by lineage. For example, in heterobranchs, *Berghia stephanieae* (Aeolidiidae) had 2049 gene family expansions and 1412 contractions, while *Aplysia californica* (Aplysiidae) showed 1513 gains and 882 losses. Their shared ancestor also underwent considerable changes (1222 expansions, 1360 contractions), pointing to a dynamic history of gene content evolution within Heterobranchia. The ampullariids *Marisa cornuarietis* and *Pomacea maculata* exhibited a moderate profile: *M. cornuarietis* had 544 expansions and 323 contractions, and *P. maculata* had 489 gains and 500 losses. In contrast, *Neverita josephinia* (Littorinimorpha) was marked by a strong contraction bias, with 7260 gene families lost and only 274 expanded **(Figure 5)**.

Taken together, these results highlight *Melo melo* as a representative of the Volutidae family with a particularly strong signal of gene family expansion relative to both other neogastropods and more distantly related caenogastropods. The high turnover observed across neogastropods further emphasizes the dynamic nature of gene family evolution within this group, with *M. melo* standing out for its prominent gene expansion profile.

### Gene Family Expansions and Adaptive Significance

#### Predatory Lifestyle and Neurogenomic Adaptations

##### Peptidase Gene Expansion and Divergence in *Melo melo*

As an active benthic predator, *Melo melo* relies on effective prey immobilization and digestion (*89*), processes often underpinned by proteolytic enzymes. To explore this, we analyzed the repertoire and copy number variation of seven peptidase gene families known for their roles in molluscan carnivory—namely Peptidase_M1, M13, M14, M28 (metallopeptidases), and Peptidase_C1, C14 (cysteine peptidases). These gene families contribute to diverse enzymatic activities, including collagen degradation, host tissue lysis, and intracellular proteolysis (*90, 91*).

*Melo melo* exhibits notably high gene counts across several of these families, with 27 copies of Peptidase_C1 and 18 of Peptidase_M14—exceeding those reported in other carnivorous neogastropods such as *Kalloconus canariensis* (Peptidase_C1: 8; Peptidase_M14: 9) and *Monoplex corrugatus* (Peptidase_C1: 10; M14: 11) **(Figure 6a, Supplementary Table 4)**. These patterns suggest that *M. melo* has undergone notable copy number expansion in digestion-associated proteases, relative to other neogastropods. Notably, orthology-based comparisons revealed that *M. melo* shares relatively few one-to-one peptidase orthologs with these taxa **(Figure 6b)**, suggesting gene diversification despite similar feeding ecologies.

**Figure 6.**
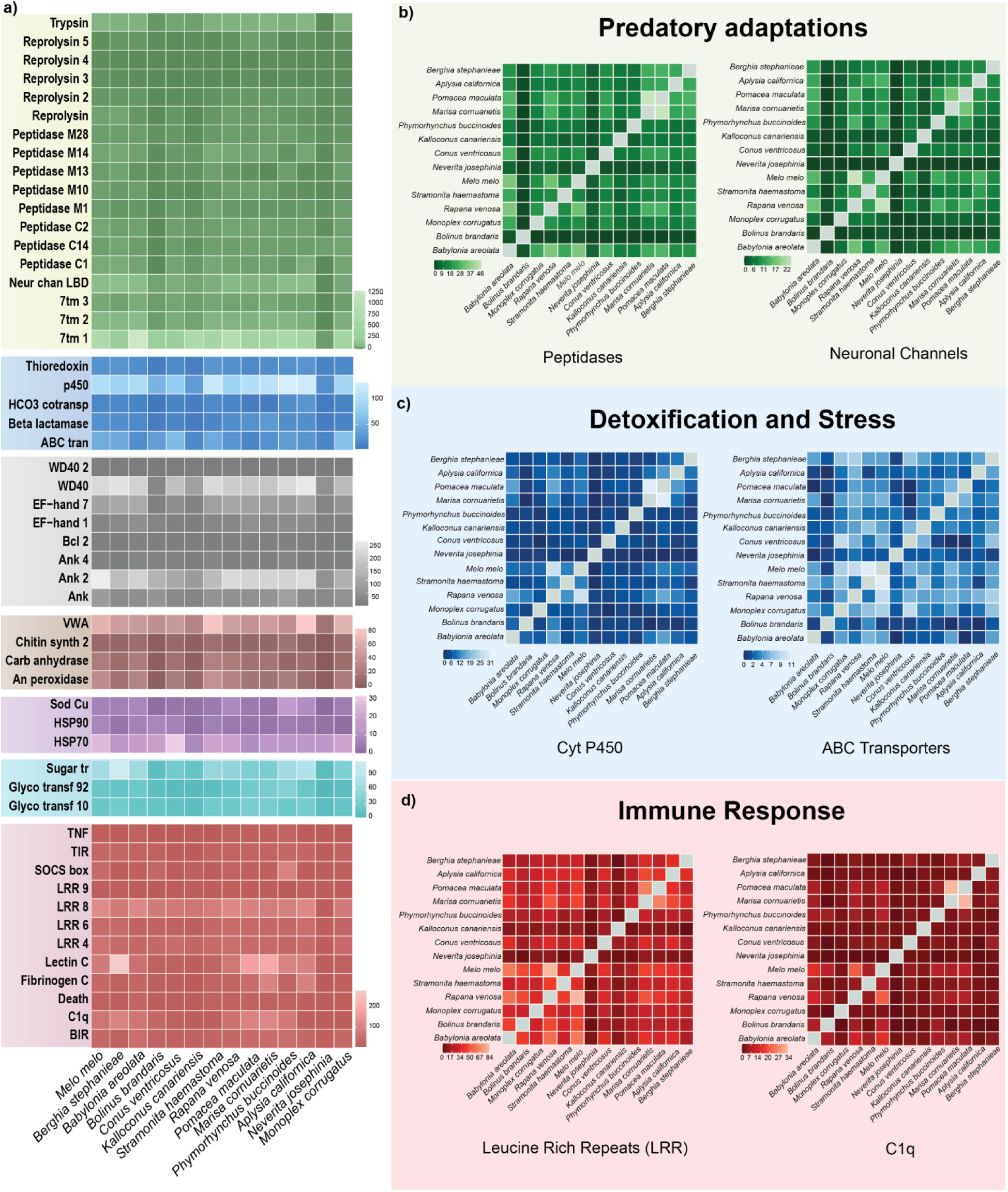
Comparative Gene family expansions and comparative orthology patterns across gastropods. **(a)** Heatmap showing total gene counts for selected gene families across 14 gastropod species. Families are grouped by biological function, including peptidases, neuronal channels, detoxification enzymes (e.g., Cytochrome P450, ABC transporters), immune-related domains (e.g., C1q, LRR, TIR), and other stress or structural proteins. Color intensity reflects gene copy number per species. **(b–d)** Orthogroup overlap plots for selected functional themes, highlighting reduced one-to-one orthology between *Melo melo* and several other gastropods. **(b)** Predatory adaptations (peptidases, neuronal channels); **(c)** Detoxification and stress (Cytochrome P450s, ABC transporters); **(d)** Immune response (Leucine-Rich Repeats, C1q). Color bars indicate the number of shared genes per species pair.

GO and KEGG enrichment analyses of *M. melo*’s predicted proteome further support this trend, revealing overrepresentation of proteolysis-related categories including “serine-type endopeptidase activity,” “lysosomal pathway,” and “proteasome complex” **(Figure 7a, b)**. This concordance between elevated copy numbers and functional enrichment indicates a broad remodeling of the proteolytic machinery in *M. melo*, potentially reflecting adaptations for prey processing in soft-sediment tropical habitats. Together, these results indicate that gene family expansion—rather than conserved one-to-one retention—may underlie the proliferation and functional diversification of peptidase repertoires in *M. melo*, contributing to its capacity for prey capture and enzymatic digestion.

**Figure 7.**
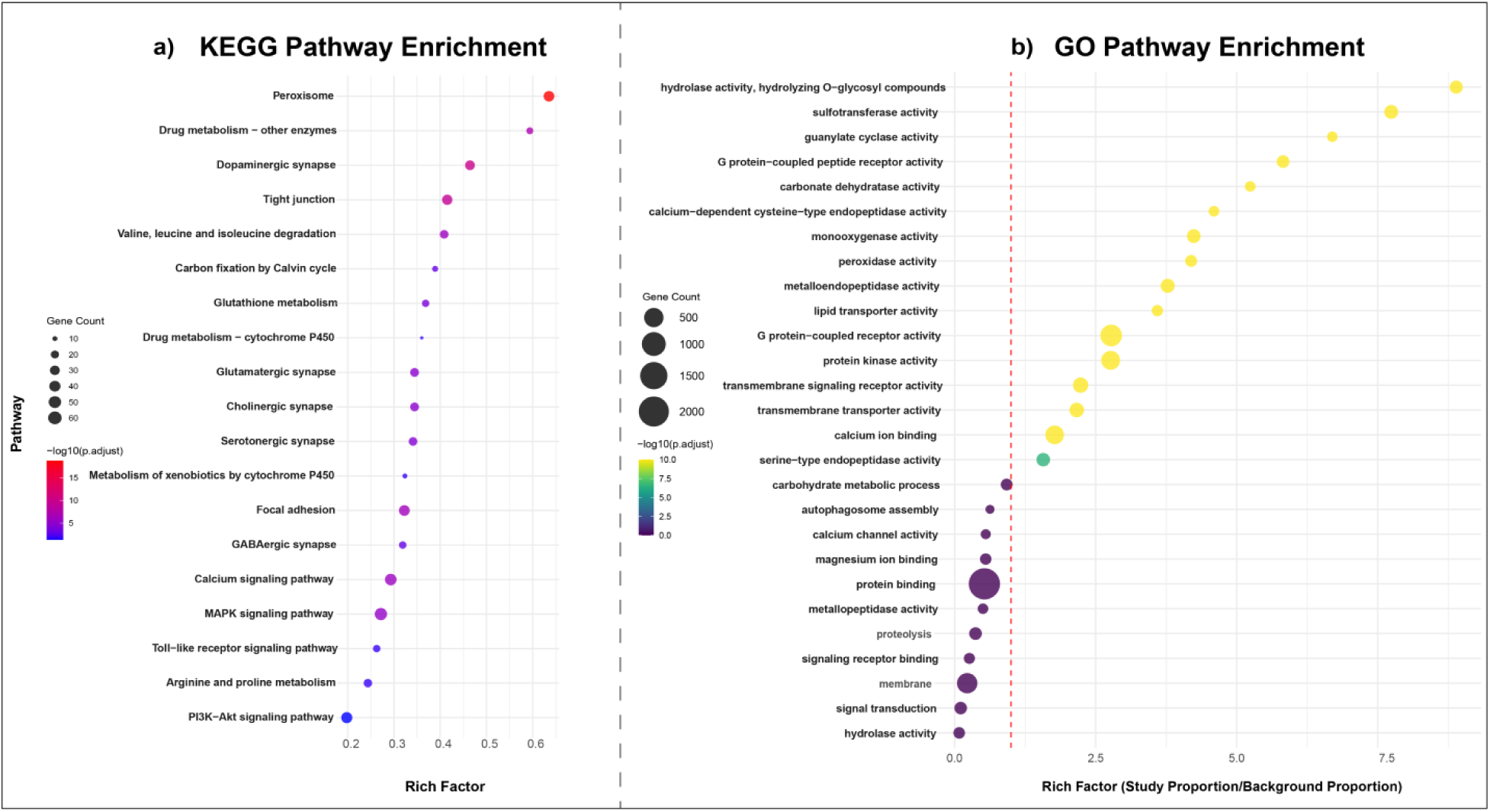
Functional enrichment of expanded gene families in *Melo melo*. (a) KEGG pathway enrichment reveals significant overrepresentation of genes involved in detoxification, synaptic signaling, and xenobiotic metabolism. (b) GO enrichment highlights dominant roles for transmembrane transport, receptor activity, and peptidase-related functions. Bubble size reflects gene count; color bars indicate adjusted p-values.

#### Conotoxin-like Genes and Potential Neurotoxic Components

In addition to proteolytic enzymes, many carnivore gastropods employ neuroactive peptides to facilitate prey immobilization (*92–95*). Among the well-characterized of these are conotoxins— small, disulfide-rich peptides predominantly studied in cone snails (*Conus* spp.)—which target a range of ion channels, neurotransmitter receptors, and synaptic proteins in prey nervous systems (*96, 97*). To investigate whether similar components might be present in *Melo melo*, we examined its genome for conotoxin-like genes. A total of 29 putative genes were identified, each encoding peptides with characteristic cysteine frameworks typical of conotoxins **(Supplementary Table 5)**. Although originally thought to be unique to conoideans, recent studies have reported low-copy homologs in other neogastropods such as *Monoplex corrugatus* and *Stramonita haemastoma* (*28*). The presence of such genes in *M. melo*, as a volutid representative, suggests that conotoxin-like sequences may have a broader distribution across predatory neogastropods than previously recognized (*98*). However, without direct evidence of their expression or functional role, the biological significance of these genes remains to be clarified. Nevertheless, these findings provide a basis for future investigation into the molecular mechanisms underlying prey capture and neurogenomic adaptation in *M. melo*.

#### Diversification of Neuronal Channel-Associated Genes

Rapid neural signaling and sensory integration are key requirements for efficient predation in gastropods (*99*). To assess this dimension in Melo melo, we focused on Pfam families related to ion channels, which include receptors involved in synaptic transmission, sensory perception, and motor coordination. For instance, the genome of *M. melo* encodes 91 Neur_chan_LBD-containing genes—a substantially higher count than in other neogastropods analyzed **(Figure 6a, Supplementary Table 4)**, including *Conus ventricosus* (38), *Kalloconus canariensis* (21), and *Monoplex corrugatus* (23) **(Figure 6a, b, Supplementary Table 4)**.

Complementing this pattern, *M. melo* also exhibits a notable abundance of G protein-coupled receptors (GPCRs) bearing the 7tm_1 domain, characteristic of chemosensory and neuro- signaling receptors. A total of 704 7tm_1-domain genes were identified in *M. melo* **(Figure 6a, Supplementary Table 4)**, a count exceeding that of several related taxa such as *Stramonita haemastoma* (561), *Phymorhynchus buccinoides* (609), and *Kalloconus canariensis* (451), though slightly lower than *Rapana venosa* (986) and markedly below *Babylonia areolata* (1302). These differences in copy number reflect varying levels of receptor diversification across neogastropods, potentially linked to habitat or prey detection strategies. KEGG pathway analysis of *M. melo* genes further showed enrichment in multiple neurotransmission-related categories, including cholinergic, glutamatergic, GABAergic, and dopaminergic synapses **(Figure 7a)**, supporting the functional relevance of the observed expansions.

### Expanded Detoxification Machinery in a Benthic Context

As a large gastropod inhabiting silty and sediment-rich coastal substrates, *Melo melo* likely encounters elevated levels of environmental toxins and microbial byproducts (*31, 100, 101*). To assess potential genomic adaptations for coping with such abiotic stressors, we examined two major gene families involved in xenobiotic processing: cytochrome P450 monooxygenases (P450s) and ATP-binding cassette (ABC) transporters.

A total of 104 P450 genes were annotated in *M. melo*, placing it among the higher end of neogastropods analyzed **(Figure 6a, Supplementary Table 4)**. This repertoire exceeds that of *Conus ventricosus* (75), *Kalloconus canariensis* (31), and *Bolinus brandaris* (53), and is comparable to sediment-dwelling carnivores such as *Stramonita haemastoma* (124) and *Rapana venosa* (86) **(Figure 6a,c Supplementary Table 4)**. Although P450 enzymes are broadly conserved in molluscs, their copy number variation across species suggests differential demands for oxidative metabolism in distinct habitats. In *M. melo*, several P450 subfamilies annotated via Pfam (e.g., PF00067) were associated with oxidation–reduction processes and metabolic detoxification **(Supplementary Table 6)**. KEGG analysis further supports this role, revealing overrepresentation of pathways related to xenobiotic degradation, drug metabolism, and retinol biosynthesis **(Figure 7a)**.

ABC transporters, which mediate the efflux of a wide range of toxic compounds, among others, were similarly expanded in *M. melo*, with 47 genes **(Figure 6a,c, Supplementary Table 4)**. This is higher than counts observed in freshwater species such as *Pomacea maculata* (11) and *Marisa cornuarietis* (12), and on par with marine carnivores like *S. haemastoma* (45) and *C. ventricosus* (60) **(Figure 6a, Supplementary Table 4)**. Functional annotation indicated enrichment in ATPase activity and transmembrane transport (GO terms) **(Figure 7b)**, reinforcing their likely role in cellular detoxification. While some one-to-one orthologs were conserved between *M. melo* and other neogastropods **(Figure 6c)**, many ABC genes showed no identifiable orthology in *M. melo*, hinting at repertoire diversification within Volutidae.

### Innate Immunity Gene Family Variation in *Melo melo*

The *Melo melo* genome encodes a broad repertoire of innate immune genes, most prominently across leucine-rich repeat (LRR) and C1q domain-containing families—two core components of the invertebrate immune recognition system (*102–104*). Among LRR-containing proteins, M. melo encodes a total of 235 genes, indicating a considerable expansion **(Figure 6a, Supplementary Table 4)**. These counts are comparable to those seen in other marine neogastropods such as *Rapana venosa* and *Stramonita haemastoma* **(Figure 6a, Supplementary Table 4)**, and align with general trends observed in molluscs known for complex immune repertoires (*105*).

The functional diversity of these LRRs was supported by Pfam annotations (e.g., PF13855, PF12799, PF13516, PF14580), many of which are linked to innate immune signaling and developmental roles **(Supplementary Table 6)**. Similarly, 64 genes encoding C1q domains were detected in *M. melo*, exceeding those in several neogastropods (*C. ventricosus*: 13; *K. canariensis*: 9) **(Figure 6a, Supplementary Table 4)**, although moderately lower than in select non-neogastropods such as *Pomacea maculata* and *Berghia stephanieae* **(Figure 6a, Supplementary Table 4)**. Gene Ontology enrichment highlighted immune-associated terms including complement activation and pathogen recognition **(Figure 7b)**. Orthology-based comparisons with other neogastropods revealed few one-to-one matches in either gene family, suggesting that much of the immune repertoire in *M. melo* consists of divergent or copy-variable paralogs **(Figure 6d)**. These patterns indicate a structurally diverse immune gene set, potentially shaped by the species’ environmental exposures and microbial context.

### Biomineralization and Shell Formation Genes

*Melo melo* produces large, non-nacreous pearls that differ structurally from the nacreous types seen in pteriid oysters(*106*) **(Zhang et al., 2012)**. To explore the genomic basis of this trait, we examined gene families implicated in molluscan shell and pearl formation. The genome encodes an expanded set of genes related to matrix secretion, cross-linking and mineral deposition: An-peroxidases (39 genes) and von willebrand factor A (VWA) domain-containing genes (68), alongside moderate representation of carbonic anhydrases and chitin synthases **(Figure 6a, Supplementary Table 4)**. In addition, the nacre-related genes such as *nacrein*, *aspein*, and *shematrin* were not found in annotated genes of *M. melo* **(Supplementary 6)**, consistent with *M. melo*’s non-nacreous shell phenotype (*107*). KEGG enrichment recovered pathways associated with calcium signaling, oxidative metabolism, and extracellular matrix organization **(Figure 7a)**, all relevant to shell and pearl biomineralization processes. Collectively, these findings indicate that *M. melo* likely employs a distinct molecular toolkit for shell formation, relying on conserved yet compositionally modified gene families within Volutidae.

## Conclusion

The high-quality genome of *Melo melo* presented here offers a rare and valuable genomic window into the early diversification of Volutidae within Neogastropoda. As the first long-read-based assembly from this family, the *M. melo* genome bridges a significant phylogenetic gap and provides a critical foundation for understanding both the ancestral genomic architecture and the functional innovations that have likely shaped predatory and benthic adaptations in early-branching neogastropods.

Our integrative analysis reveals extensive gene family expansions related to prey digestion, venom production, neuronal signal transduction, detoxification, and innate immunity. These expansions likely underpin the complex behavioral and physiological traits associated with *M. melo*’s active predatory lifestyle and its adaptation to silty, microbe-rich benthic habitats. Notably the presence of a diverse set of conotoxin-like genes and a broad repertoire of peptidases and neuronal channels, suggesting that *M. melo*—and perhaps Volutidae more broadly—harbor a distinct molecular toolkit that enables effective prey capture, immobilization, and processing. Furthermore, the pronounced expansion of detoxification and immune-related genes reflects the environmental challenges associated with sediment-dwelling life. In the context of biomineralization, *M. melo*’s genomic profile provides novel insights into the production of its large, non-nacreous pearls. The absence of typical nacre-specific genes, combined with the enrichment of alternative matrix-related gene families and calcium-regulating pathways, points to a modified but functionally robust shell formation mechanism within Volutidae. This divergence from the nacreous model highlights the potential for lineage-specific variation in molluscan biomineralization strategies.

Together, these findings establish *Melo melo* as a pivotal reference for investigating genomic and functional evolution across Volutidae and early neogastropods. This study highlights the importance of future work aimed at understanding how these genes function across different tissues and life stages. Approaches such as tissue-specific transcriptomics and proteomics will be key to linking gene expansions with ecological traits in *Melo melo*. With the future availability of more volutid genomes, these findings can serve as a baseline for exploring the genomic underpinnings of evolutionary adaptation in the Neogastropod tree of life.

## Supplementary Data

The data underlying this article are available within the article and its online supplementary material. All FASTA sequences, Pfam annotation files, genome statistics, repeat counts, and the taxa list are also accessible at https://doi.org/10.5281/zenodo.17490824.

## Data Availability

All raw sequencing data generated in this study have been deposited in the NCBI Sequence Read Archive (SRA). The genome sequencing reads are available under accession SRR30898481, and the transcriptome sequencing reads under SRR30899298. Both datasets are linked to BioProject **PRJNA1153738** and BioSample **SAMN43402451**. The assembled genome has been submitted to NCBI under the accession **JBGTYI000000000**.

## References

1. R. L. Cunha, C. Grande, R. Zardoya, Neogastropod phylogenetic relationships based on entire mitochondrial genomes. BMC Evolutionary Biology 9, 210 (2009).

2. A. Wanninger, T. Wollesen, The evolution of molluscs. Biological Reviews 94, 102–115 (2019).

3. W. F. Ponder, D. R. Lindberg, J. M. Ponder, Biology and evolution of the mollusca, volume 1 (crc Press, 2019).

4. A. E. Fedosov, P. Zaharias, T. Lemarcis, M. V. Modica, M. Holford, M. Oliverio, Y. I. Kantor, N. Puillandre, Phylogenomics of Neogastropoda: the backbone hidden in the bush. Systematic Biology 73, 521–531 (2024).

5. S. Dominici, The Revolution of Small Snails and the Early Modern Evolutionary Fauna. Diversity (14242818) **17**, (2025).

6. G. Zancolli, M. V. Modica, N. Puillandre, Y. Kantor, A. Barua, G. Campli, M. Robinson-Rechavi, Redistribution of ancestral functions underlies the evolution of venom production in marine predatory snails. Molecular biology and evolution 42, msaf095 (2025).

7. M. V. Modica, F. Lombardo, P. Franchini, M. Oliverio, The venomous cocktail of the vampire snail Colubraria reticulata (Mollusca, Gastropoda). BMC genomics 16, 441 (2015).

8. B. M. Olivera, A. Fedosov, J. S. Imperial, Y. Kantor, "Physiology of envenomation by conoidean gastropods" in Physiology of molluscs (Apple Academic Press, 2017), pp. 169–204.

9. L. R. Page, Metamorphic remodeling of a planktotrophic larva to produce the predatory feeding system of a cone snail (Mollusca, Neogastropoda). The Biological Bulletin 221, 176–188 (2011).

10. E. Vortsepneva, A. Mikhlina, Y. Kantor, Main patterns of radula formation and ontogeny in Gastropoda. Journal of Morphology 284, e21538 (2023).

11. Z.-L. Yu, M.-J. Yang, H. Song, T. Zhang, X.-T. Yuan, Gastropod chemoreception behaviors—Mechanisms underlying the perception and location of targets and implications for shellfish fishery development in aquatic environments. Frontiers in Marine Science 9, 1042962 (2023).

12. D. R. Lindberg, J. D. Sigwart, What is the molluscan osphradium? A reconsideration of homology. Zoologischer Anzeiger-A Journal of Comparative Zoology 256, 14–21 (2015).

13. T. Jd, Food specialization and the evolution of predatory prosobranch gastropods. Palaeontology 23, 375–409 (1980).

14. P. R. Gutthavilli, A. M. Bharne, K. Marimuthu, G. Thiruchitrambalam, Unveiling the enigmatic cone snails along the coastal environments of the South Andaman Islands: diversity, distribution and their habitat preference. Frontiers in Marine Science 11, 1477472 (2024).

15. E. M. Montgomery, J.-F. Hamel, A. Mercier, The deep-sea neogastropod Buccinum scalariforme: Reproduction, development and growth. Deep Sea Research Part I: Oceanographic Research Papers 119, 24–33 (2017).

16. Y. I. Kantor, N. Puillandre, K. Fraussen, A. Fedosov, P. Bouchet, Deep-water Buccinidae (Gastropoda: Neogastropoda) from sunken wood, vents and seeps: molecular phylogeny and taxonomy. Journal of the Marine Biological Association of the United Kingdom 93, 2177–2195 (2013).

17. F. Criscione, A. Hallan, N. Puillandre, A. Fedosov, Where the snails have no name: a molecular phylogeny of Raphitomidae (Neogastropoda: Conoidea) uncovers vast unexplored diversity in the deep seas of temperate southern and eastern Australia. Zoological Journal of the Linnean Society 191, 961–1000 (2021).

18. C. J. Plante, K. M. Hill-Spanik, R. Emerson, Feeding effects of the keystone deposit feeder Ilyanassa obsoleta (Neogastropoda, Gastropoda) on sedimentary diatoms. Journal of Phycology 59, 590–602 (2023).

19. E. E. Strong, Refining molluscan characters: morphology, character coding and a phylogeny of the Caenogastropoda. Zoological Journal of the Linnean Society 137, 447–554 (2003).

20. R. E. Golding, W. F. Ponder, Homology and morphology of the neogastropod valve of Leiblein (Gastropoda: Caenogastropoda). Zoomorphology 129, 81–91 (2010).

21. Y. I. Kantor, E. E. Strong, N. Puillandre, A new lineage of Conoidea (Gastropoda: Neogastropoda) revealed by morphological and molecular data. Journal of Molluscan Studies 78, 246–255 (2012).

22. A. Nützel, Larval ecology and morphology in fossil gastropods. Palaeontology 57, 479–503 (2014).

23. Z. Ratibou, N. Inguimbert, S. Dutertre, Predatory and defensive strategies in cone snails. Toxins 16, 94 (2024).

24. M. V. Modica, J. Gorson, A. E. Fedosov, G. Malcolm, Y. Terryn, N. Puillandre, M. Holford, Macroevolutionary analyses suggest that environmental factors, not venom apparatus, play key role in terebridae marine snail diversification. Systematic Biology 69, 413–430 (2020).

25. P. E. Bourdeau, Intraspecific trait cospecialization of constitutive and inducible morphological defences in a marine snail from habitats with different predation risk. Journal of Animal Ecology 81, 849–858 (2012).

26. P. Bouchet, J.-P. Rocroi, B. Hausdorf, A. Kaim, Y. Kano, A. Nützel, P. Parkhaev, M. Schrödl, E. E. Strong, Revised classification, nomenclator and typification of gastropod and monoplacophoran families. Malacologia 61, 1–526 (2017).

27. J. R. Pardos-Blas, I. Irisarri, S. Abalde, C. M. Afonso, M. J. Tenorio, R. Zardoya, The genome of the venomous snail Lautoconus ventricosus sheds light on the origin of conotoxin diversity. Gigascience 10, giab037 (2021).

28. S. Farhat, M. V. Modica, N. Puillandre, Whole genome duplication and gene evolution in the hyperdiverse venomous gastropods. Molecular Biology and Evolution 40, msad171 (2023).

29. R. Nath, B. Panda, S. Rakesh, A. Krishnan, Lineage-Specific Class-A GPCR Dynamics Reflect Diverse Chemosensory Adaptations in Lophotrochozoa. Molecular Biology and Evolution 42, msaf042 (2025).

30. G. J. Vermeij, V. M. Watson-Zink, Terrestrialization in gastropods: lineages, ecological constraints and comparisons with other animals. Biological Journal of the Linnean Society 136, 393–404 (2022).

31. L. Aristide, R. Fernández, Genomic insights into mollusk terrestrialization: parallel and convergent gene family expansions as key facilitators in out-of-the-sea transitions. Genome Biology and Evolution 15, evad176 (2023).

32. C. Liu, Y. Ren, Z. Li, Q. Hu, L. Yin, H. Wang, X. Qiao, Y. Zhang, L. Xing, Y. Xi, Giant African snail genomes provide insights into molluscan whole-genome duplication and aquatic–terrestrial transition. Molecular Ecology Resources 21, 478–494 (2021).

33. J. Sun, C. Chen, N. Miyamoto, R. Li, J. D. Sigwart, T. Xu, Y. Sun, W. C. Wong, J. C. Ip, W. Zhang, The Scaly-foot Snail genome and implications for the origins of biomineralised armour. Nature communications 11, 1657 (2020).

34. X. Zeng, Y. Zhang, L. Meng, G. Fan, J. Bai, J. Chen, Y. Song, I. Seim, C. Wang, Z. Shao, Genome sequencing of deep-sea hydrothermal vent snails reveals adaptions to extreme environments. Gigascience 9, giaa139 (2020).

35. Z. Liu, Y. Huang, H. Chen, C. Liu, M. Wang, C. Bian, L. Wang, L. Song, Chromosome-level genome assembly of the deep-sea snail Phymorhynchus buccinoides provides insights into the adaptation to the cold seep habitat. BMC genomics 24, 679 (2023).

36. S. Espino, M. Watkins, R. Probst, T. L. Koch, K. Chase, J. Imperial, S. D. Robinson, P. Flórez Salcedo, D. Taylor, J. Gajewiak, χ-Conotoxins are an evolutionary innovation of mollusk-hunting cone snails as a counter-adaptation to prey defense. Molecular Biology and Evolution 41, msae226 (2024).

37. G. Chiappa, G. Fassio, M. V. Modica, M. Oliverio, Potential Ancestral Conoidean Toxins in the Venom Cocktail of the Carnivorous Snail Raphitoma purpurea (Montagu, 1803)(Neogastropoda: Raphitomidae). Toxins 16, 348 (2024).

38. M. Shen, G. Di, M. Li, J. Fu, Q. Dai, X. Miao, M. Huang, W. You, C. Ke, Proteomics studies on the three larval stages of development and metamorphosis of Babylonia areolata. Scientific Reports 8, 6269 (2018).

39. Y. Zou, J. Fu, Y. Liang, X. Luo, M. Shen, M. Huang, Y. Chen, W. You, C. Ke, Chromosome-level genome assembly of the ivory shell Babylonia areolata. Scientific Data 11, 1201 (2024).

40. H. Song, Z. Li, M. Yang, P. Shi, Z. Yu, Z. Hu, C. Zhou, P. Hu, T. Zhang, Chromosome-level genome assembly of the caenogastropod snail Rapana venosa. Scientific Data 10, 539 (2023).

41. C. Peng, Y. Huang, C. Bian, J. Li, J. Liu, K. Zhang, X. You, Z. Lin, Y. He, J. Chen, The first Conus genome assembly reveals a primary genetic central dogma of conopeptides in C. betulinus. Cell discovery 7, 11 (2021).

42. S. Tracey, Mollusca: gastropoda. The fossil record *2*, (1993).

43. M. Harasewych, M. Sei, A. Oleinik, J. E. Uribe, The complete mitochondrial genome of Voluta musica Linnaeus, 1758 (Neogastropoda: Volutidae: Volutinae). The Nautilus 138, 1-7 (2024).

44. M. Ojeda, F. Arrighetti, J. Giménez, Morphology and cyclic activity of the digestive gland of Zidona dufresnei (Caenogastropoda: Volutidae). Malacologia 58, 157–165 (2015).

45. M. Harasewych, Tractolira germonae, a new abyssal Antarctic volute. The Nautilus 101, 3–8 (1987).

46. G. Bigatti, C. Sanchez Antelo, P. Miloslavich, P. E. Penchaszadeh, Feeding behavior of Adelomelon ancilla (Ligfoot, 1786): A predatory neogastropod (Gastropoda: Volutidae) in Patagonian benthic communities. (2009).

47. B. Morton, The diet and prey capture mechanism of Melo melo (Prosobranchia: Volutidae). Journal of Molluscan Studies 52, 156–160 (1986).

48. H. Cheng, G. T. Concepcion, X. Feng, H. Zhang, H. Li, Haplotype-resolved de novo assembly using phased assembly graphs with hifiasm. Nature methods 18, 170–175 (2021).

49. M. Alonge, L. Lebeigle, M. Kirsche, K. Jenike, S. Ou, S. Aganezov, X. Wang, Z. B. Lippman, M. C. Schatz, S. Soyk, Automated assembly scaffolding using RagTag elevates a new tomato system for high-throughput genome editing. Genome biology 23, 258 (2022).

50. R. Challis, E. Richards, J. Rajan, G. Cochrane, M. Blaxter, BlobToolKit–interactive quality assessment of genome assemblies. G3: Genes, Genomes, Genetics 10, 1361–1374 (2020).

51. M. Uliano-Silva, J. G. R. Ferreira, K. Krasheninnikova, G. Formenti, L. Abueg, J. Torrance, E. W. Myers, R. Durbin, M. Blaxter, MitoHiFi: a python pipeline for mitochondrial genome assembly from PacBio high fidelity reads. BMC bioinformatics 24, 288 (2023).

52. M. Bernt, A. Donath, F. Jühling, F. Externbrink, C. Florentz, G. Fritzsch, J. Pütz, M. Middendorf, P. F. Stadler, MITOS: improved de novo metazoan mitochondrial genome annotation. Molecular phylogenetics and evolution 69, 313–319 (2013).

53. J. R. Grant, E. Enns, E. Marinier, A. Mandal, E. K. Herman, C.-y. Chen, M. Graham, G. Van Domselaar, P. Stothard, Proksee: in-depth characterization and visualization of bacterial genomes. Nucleic acids research 51, W484–W492 (2023).

54. F. A. Simão, R. M. Waterhouse, P. Ioannidis, E. V. Kriventseva, E. M. Zdobnov, BUSCO: assessing genome assembly and annotation completeness with single-copy orthologs. Bioinformatics 31, 3210–3212 (2015).

55. A. Gurevich, V. Saveliev, N. Vyahhi, G. Tesler, QUAST: quality assessment tool for genome assemblies. Bioinformatics 29, 1072–1075 (2013).

56. S. W. Wingett, S. Andrews, FastQ Screen: A tool for multi-genome mapping and quality control. F1000Research **7**, 1338 (2018).

57. W. Bao, K. K. Kojima, O. Kohany, Repbase Update, a database of repetitive elements in eukaryotic genomes. Mobile Dna 6, 11 (2015).

58. J. M. Flynn, R. Hubley, C. Goubert, J. Rosen, A. G. Clark, C. Feschotte, A. F. Smit, RepeatModeler2 for automated genomic discovery of transposable element families. Proceedings of the National Academy of Sciences 117, 9451–9457 (2020).

59. T. Brůna, K. J. Hoff, A. Lomsadze, M. Stanke, M. Borodovsky, BRAKER2: automatic eukaryotic genome annotation with GeneMark-EP+ and AUGUSTUS supported by a protein database. NAR genomics and bioinformatics 3, lqaa108 (2021).

60. H. Li, Minimap2: pairwise alignment for nucleotide sequences. Bioinformatics 34, 3094–3100 (2018).

61. T. Brůna, A. Lomsadze, M. Borodovsky, GeneMark-ETP significantly improves the accuracy of automatic annotation of large eukaryotic genomes. Genome Research 34, 757–768 (2024).

62. L. Fu, B. Niu, Z. Zhu, S. Wu, W. Li, CD-HIT: accelerated for clustering the next-generation sequencing data. Bioinformatics 28, 3150–3152 (2012).

63. X. Li, W. Chen, D. Zhangsun, S. Luo, Diversity of conopeptides and their precursor genes of Conus litteratus. Marine drugs 18, 464 (2020).

64. M. A. Phuong, G. N. Mahardika, Targeted sequencing of venom genes from cone snail genomes improves understanding of conotoxin molecular evolution. Molecular biology and evolution 35, 1210–1224 (2018).

65. S. R. Eddy, Accelerated profile HMM searches. PLoS computational biology 7, e1002195 (2011).

66. Q. Kaas, R. Yu, A.-H. Jin, S. Dutertre, D. J. Craik, ConoServer: updated content, knowledge, and discovery tools in the conopeptide database. Nucleic acids research 40, D325–D330 (2012).

67. B. J. Haas, A. Papanicolaou, M. Yassour, M. Grabherr, P. D. Blood, J. Bowden, M. B. Couger, D. Eccles, B. Li, M. Lieber, De novo transcript sequence reconstruction from RNA-seq using the Trinity platform for reference generation and analysis. Nature protocols 8, 1494–1512 (2013).

68. K. M. Kocot, T. H. Struck, J. Merkel, D. S. Waits, C. Todt, P. M. Brannock, D. A. Weese, J. T. Cannon, L. L. Moroz, B. Lieb, Phylogenomics of Lophotrochozoa with consideration of systematic error. Systematic biology 66, 256–282 (2017).

69. X. Liu, J. D. Sigwart, J. Sun, Phylogenomic analyses shed light on the relationships of chiton superfamilies and shell-eye evolution. Marine Life Science & Technology 5, 525–537 (2023).

70. D. M. Emms, S. Kelly, OrthoFinder: phylogenetic orthology inference for comparative genomics. Genome biology 20, 238 (2019).

71. K. Katoh, K. Misawa, K. i. Kuma, T. Miyata, MAFFT: a novel method for rapid multiple sequence alignment based on fast Fourier transform. Nucleic acids research 30, 3059–3066 (2002).

72. A. Di Franco, R. Poujol, D. Baurain, H. Philippe, Evaluating the usefulness of alignment filtering methods to reduce the impact of errors on evolutionary inferences. BMC Evolutionary Biology 19, 21 (2019).

73. A. Criscuolo, S. Gribaldo, BMGE (Block Mapping and Gathering with Entropy): a new software for selection of phylogenetic informative regions from multiple sequence alignments. BMC evolutionary biology 10, 210 (2010).

74. F. Thalén, PhyloPyPruner: tree-based orthology inference for phylogenomics with new methods for identifying and excluding contamination. (2018).

75. B. Q. Minh, H. A. Schmidt, O. Chernomor, D. Schrempf, M. D. Woodhams, A. Von Haeseler, R. Lanfear, IQ-TREE 2: new models and efficient methods for phylogenetic inference in the genomic era. Molecular biology and evolution 37, 1530–1534 (2020).

76. M. R. Ford, S. V. Vollmer, G. C. Trussell, Annotated genome of the Atlantic dog whelk, Nucella lapillus. G3: Genes, Genomes, Genetics 15, jkaf182 (2025).

77. J. Jurka, V. V. Kapitonov, O. Kohany, M. V. Jurka, Repetitive sequences in complex genomes: structure and evolution. Annu. Rev. Genomics Hum. Genet. 8, 241–259 (2007).

78. J. A. Baeza, M. T. González, J. D. Sigwart, C. Greve, S. Pirro, Insights into the genome of the ‘Loco’Concholepas concholepas (Gastropoda: Muricidae) from low-coverage short-read sequencing: genome size, ploidy, transposable elements, nuclear RNA gene operon, mitochondrial genome, and phylogenetic placement in the family Muricidae. BMC genomics 25, 77 (2024).

79. Z. Chen, Ö. Doğan, N. Guiglielmoni, A. Guichard, M. Schrödl, Pulmonate slug evolution is reflected in the de novo genome of Arion vulgaris Moquin-Tandon, 1855. Scientific reports **12**, 14226 (2022).

80. A. Gomes-dos-Santos, M. Lopes-Lima, L. F. C. Castro, E. Froufe, Molluscan genomics: the road so far and the way forward. Hydrobiologia 847, 1705–1726 (2020).

81. H. Wang, X. He, C. Chen, K. Gao, Y. Dai, J. Sun, New insights into the phylogeny of Neogastropoda aided by draft genome sequencing of a volutid snail. Zoologica Scripta 53, 805–817 (2024).

82. A. E. Fedosov, M. Caballer Gutierrez, B. Buge, P. V. Sorokin, N. Puillandre, P. Bouchet, Mapping the missing branch on the neogastropod tree of life: molecular phylogeny of marginelliform gastropods. Journal of Molluscan Studies 85, 439–451 (2019).

83. T. Lemarcis, A. E. Fedosov, Y. I. Kantor, J. Abdelkrim, P. Zaharias, N. Puillandre, Neogastropod (Mollusca, Gastropoda) phylogeny: a step forward with mitogenomes. Zoologica Scripta 51, 550–561 (2022).

84. L. Du, S. Cai, J. Liu, R. Liu, H. Zhang, The complete mitochondrial genome of a cold seep gastropod Phymorhynchus buccinoides (Neogastropoda: Conoidea: Raphitomidae). PLoS One 15, e0242541 (2020).

85. A. Barco, M. Claremont, D. Reid, R. Houart, P. Bouchet, S. Williams, C. Cruaud, A. Couloux, M. Oliverio, A molecular phylogenetic framework for the Muricidae, a diverse family of carnivorous gastropods. Molecular Phylogenetics and Evolution 56, 1025–1039 (2010).

86. F. Li, W. Li, Y. Zhang, A. Wang, C. Liu, Z. Gu, Y. Yang, The molecular phylogeny of Caenogastropoda (Mollusca, Gastropoda) based on mitochondrial genomes and nuclear genes. Gene 928, 148790 (2024).

87. D. K. Jacobs, D. R. Lindberg, Oxygen and evolutionary patterns in the sea: onshore/offshore trends and recent recruitment of deep-sea faunas. Proceedings of the National Academy of Sciences 95, 9396–9401 (1998).

88. F. K. Mendes, D. Vanderpool, B. Fulton, M. W. Hahn, CAFE 5 models variation in evolutionary rates among gene families. Bioinformatics 36, 5516–5518 (2020).

89. V. Teoh, I. Izzat-Irfan, A. Jaya-Ram, S. Woo, Feeding observation of the Indian volute Melo melo (Lightfoot, 1786) in captivity. International Journal of Aquatic Research and Environmental Studies 3, 101-104 (2023).

90. H. Song, Z.-L. Yu, L.-N. Sun, D.-X. Xue, T. Zhang, H.-Y. Wang, Transcriptomic analysis of differentially expressed genes during larval development of Rapana venosa by digital gene expression profiling. G3: Genes, Genomes, Genetics 6, 2181–2193 (2016).

91. A. Lu, M. Watkins, Q. Li, S. D. Robinson, G. P. Concepcion, M. Yandell, Z. Weng, B. M. Olivera, H. Safavi-Hemami, A. E. Fedosov, Transcriptomic profiling reveals extraordinary diversity of venom peptides in unexplored predatory gastropods of the genus Clavus. Genome Biology and Evolution 12, 684–700 (2020).

92. S. Himaya, A.-H. Jin, S. Dutertre, J. Giacomotto, H. Mohialdeen, I. Vetter, P. F. Alewood, R. J. Lewis, Comparative venomics reveals the complex prey capture strategy of the piscivorous cone snail Conus catus. Journal of Proteome Research 14, 4372–4381 (2015).

93. S. Himaya, A.-H. Jin, B. Hamilton, S. K. Rai, P. Alewood, R. J. Lewis, Venom duct origins of prey capture and defensive conotoxins in piscivorous Conus striatus. Scientific Reports 11, 13282 (2021).

94. S. Dutertre, A.-H. Jin, I. Vetter, B. Hamilton, K. Sunagar, V. Lavergne, V. Dutertre, B. G. Fry, A. Antunes, D. J. Venter, Evolution of separate predation-and defence-evoked venoms in carnivorous cone snails. Nature communications 5, 3521 (2014).

95. K. G. Kuznetsova, S. S. Zvonareva, R. Ziganshin, E. S. Mekhova, P. Dgebuadze, D. T. Yen, T. H. Nguyen, S. A. Moshkovskii, A. E. Fedosov, Vexitoxins: conotoxin-like venom peptides from predatory gastropods of the genus Vexillum. Proceedings of the Royal Society B 289, 20221152 (2022).

96. B. M. Olivera, P. Showers Corneli, M. Watkins, A. Fedosov, Biodiversity of cone snails and other venomous marine gastropods: evolutionary success through neuropharmacology. Annu. Rev. Anim. Biosci. 2, 487–513 (2014).

97. H. Morales Duque, S. Campos Dias, O. L. Franco, Structural and functional analyses of cone snail toxins. Marine drugs 17, 370 (2019).

98. M. A. Phuong, G. N. Mahardika, M. E. Alfaro, Dietary breadth is positively correlated with venom complexity in cone snails. BMC genomics 17, 401 (2016).

99. J. Gorson, G. Ramrattan, A. Verdes, E. M. Wright, Y. Kantor, R. Rajaram Srinivasan, R. Musunuri, D. Packer, G. Albano, W.-G. Qiu, Molecular diversity and gene evolution of the venom arsenal of terebridae predatory marine snails. Genome Biology and Evolution 7, 1761–1778 (2015).

100. J. Sun, Y. Zhang, T. Xu, Y. Zhang, H. Mu, Y. Zhang, Y. Lan, C. J. Fields, J. H. L. Hui, W. Zhang, Adaptation to deep-sea chemosynthetic environments as revealed by mussel genomes. Nature ecology & evolution 1, 0121 (2017).

101. J. Zanette, J. V. Goldstone, A. C. Bainy, J. J. Stegeman, Identification of CYP genes in Mytilus (mussel) and Crassostrea (oyster) species: first approach to the full complement of cytochrome P450 genes in bivalves. Marine environmental research 69, S1–S3 (2010).

102. J. H. Schultz, C. M. Adema, Comparative immunogenomics of molluscs. Developmental & Comparative Immunology 75, 3–15 (2017).

103. W. Wang, X. Song, L. Wang, L. Song, Pathogen-derived carbohydrate recognition in molluscs immune defense. International journal of molecular sciences 19, 721 (2018).

104. L. Zhang, L. Li, X. Guo, G. W. Litman, L. J. Dishaw, G. Zhang, Massive expansion and functional divergence of innate immune genes in a protostome. Scientific reports 5, 8693 (2015).

105. R. B. Rathinam, A. Acharya, A. J. Robina, H. Banu, G. Tripathi, The immune system of marine invertebrates: Earliest adaptation of animals. Comparative Immunology Reports 7, 200163 (2024).

106. G. Zhang, X. Fang, X. Guo, L. Li, R. Luo, F. Xu, P. Yang, L. Zhang, X. Wang, H. Qi, The oyster genome reveals stress adaptation and complexity of shell formation. Nature 490, 49–54 (2012).

107. X. Song, Z. Liu, L. Wang, L. Song, Recent advances of shell matrix proteins and cellular orchestration in marine molluscan shell biomineralization. Frontiers in Marine Science 6, 41 (2019).

