## Supplementary Figures for "Genome of the Predatory Volute *Melo melo* Provides Insights into Adaptive Gene Family Diversification in a Basal Neogastropod Lineage"

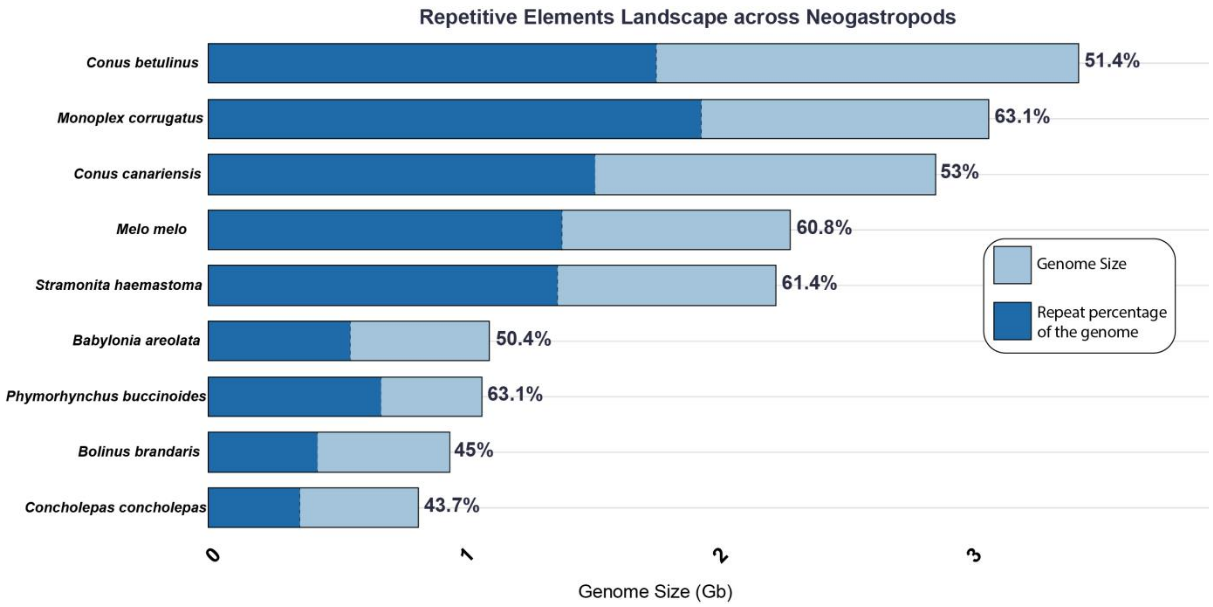


**Supplementary Fig 1:** Repetitive element content of nine representative Neogastropods have been depicted with respect to their total genomic size. The dark blue and sky blue represents the repeat percentage of the genome and genomic size respectively.

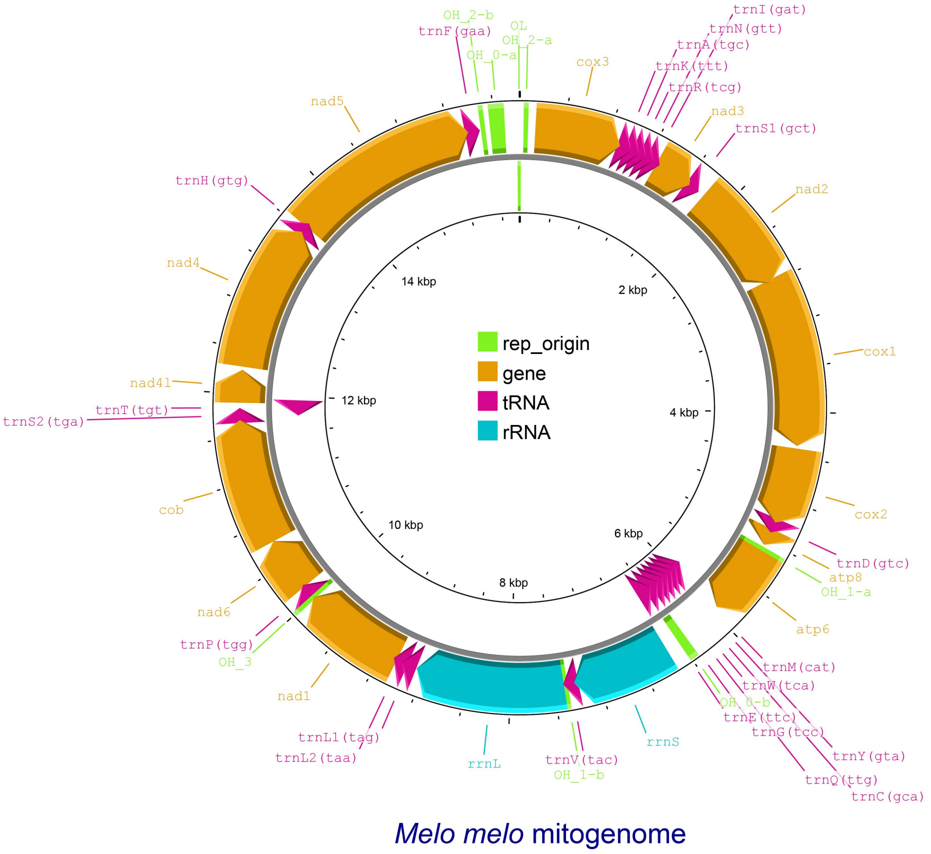


**Supplementary Fig 2:** Mitochondrial Genome of *Melo melo.*
