## Supplementary Table 1 for "Genome of the Predatory Volute *Melo melo* Provides Insights into Adaptive Gene Family Diversification in a Basal Neogastropod Lineage"

**a)**

| **HiFi Read Statistics** | |
| --- | --- |
| Total bases | 25103218299 |
| Total reads | 2852782 |
| Average read length | 8799.557 |
| A | 6966989123 |
| T | 7146403412 |
| G | 5533702120 |
| C | 5456123644 |
| GC% | 43.779 |

**b)**

| **Denovo Genome Assembly Statistics (Hifiasm)** | |
| --- | --- |
| Number of contigs | 18019 |
| Total size of contigs | 2293822963 |
| Longest contig | 5188507 |
| Shortest contig | 2089 |
| Number of contigs > 1K nt | 18019 |
| Number of contigs > 10K nt | 14788 |
| Number of contigs > 100K nt | 5093 |
| Number of contigs > 1M nt | 287 |
| Mean contig size | 127300 |
| Median contig size | 34559 |
| N50 contig length | 427368 |
| L50 contig count | 1441 |
| contig %A | 28.76 |
| contig %C | 21.22 |
| contig %G | 21.23 |
| contig %T | 28.78 |
| contig %N | 0 |

**c)**

| **Final Genome Assembly Statistics** | |
| --- | --- |
| Number of scaffolds | 13961 |
| Total size of scaffolds | 2294228763 |
| Longest scaffold | 107503566 |
| Shortest scaffold | 2089 |
| Number of scaffold > 1K nt | 13961 |
| Number of scaffold > 10K nt | 10870 |
| Number of scaffold > 100K nt | 2605 |
| Number of scaffold > 1M nt | 76 |
| Number of scaffold > 10M nt | 33 |
| Mean scaffold size | 164331 |
| Median scaffold size | 24612 |
| N50 scaffold length | 18608260 |
| L50 scaffold count | 27 |
| scaffold %A | 28.76 |
| scaffold %C | 21.22 |
| scaffold %G | 21.23 |
| scaffold %T | 28.78 |
| scaffold %N | 0 |
| GC (%) | 42.45 |

**d)**

|  |  | **number of elements** | **length occupied** | **percentage of sequence** |
| --- | --- | --- | --- | --- |
| **Retroelements** |  | 1161667 | 343768070 bp | 14.98 % |
|  | **SINEs:** | 0 | 0 bp | 0.00 % |
|  | **Penelope** | 0 | 0 bp | 0.00 % |
|  | **LINEs** | 1157271 | 339661659 bp | 14.81 % |
|  | **CRE/SLACS** | 0 | 0 bp | 0.00 % |
|  | **L2/CR1/Rex** | 459439 | 104071947 bp | 4.54 % |
|  | **R1/LOA/Jockey** | 174086 | 40859479 bp | 1.78 % |
|  | **R2/R4/NeSL** | 308 | 0 bp | 0.00 % |
|  | **RTE/Bov-B** | 480491 | 181322333 bp | 7.90 % |
|  | **L1/CIN4** | 0 | 0 bp | 0.00 % |
|  | **LTR elements:** | 4396 | 4106411 bp | 0.18 % |
|  | **BEL/Pao** | 0 | 0 bp | 0.00 % |
|  | **Ty1/Copia** | 0 | 0 bp | 0.00 % |
|  | **Gypsy/DIRS1** | 4396 | 4106411 bp | 0.18 % |
|  | **Retroviral** | 0 | 0 bp | 0.00 % |
| **DNA transposons** |  | 46417 | 13023734 bp | 0.57 % |
|  | **hobo-Activator** | 2153 | 505071 bp | 0.02 % |
|  | **Tc1-IS630-Pogo** | 9893 | 1188937 bp | 0.05 % |
|  | **En-Spm** | 0 | 0 bp | 0.00 % |
|  | **MULE-MuDR** | 0 | 0 bp | 0.00 % |
|  | **PiggyBac** | 1462 | 1407999 bp | 0.06 % |
|  | **Tourist/Harbinger** | 0 | 0 bp | 0.00 % |
|  | **Other (Mirage, P-element, Transib)** | 0 | 0 bp | 0.00 % |
|  | **Rolling-circles** | 0 | 0 bp | 0.00 % |
|  | **Unclassified:** | 3965393 | 630986843 bp | 27.50 % |
|  | **Total interspersed repeats:** | - | 987778647 bp | 43.05 % |
|  | **Small RNA:** | 19855 | 938701 bp | 0.04 % |
|  | **Satellites:** | 5 | 3848 bp | 0.00 % |
|  | **Simple repeats:** | 4111319 | 373357019 bp | 16.27 % |
|  | **Low complexity:** | 309636 | 32004380 bp | 1.39 % |

**Supplementary Table 1:** The genome statistics of *Melo melo*, a) Raw HiFi read statistics, b) Initial Denovo Genome assembly statistics of *M.melo*, c) Final genome assembly statistics for *M.melo.* d) Repetitive elements statistics of *M.melo.*
