## Supplementary Table 2 for "Genome of the Predatory Volute *Melo melo* Provides Insights into Adaptive Gene Family Diversification in a Basal Neogastropod Lineage"

| **Total Gene Counts of *M.melo*** | **29,364** |
| --- | --- |
| **Database** | **Genes Annotated** |
| PFAM | 26,502 |
| Swissprot | 24,949 |
| Interproscan | 21,055 |
| KEGG | 10,804 |

**Supplementary Table 2:** The table shows the total number of genes predicted by braker using M.melo. Functional annotation against different databases and their corresponding counts.
